# Phase-locking of hippocampal CA3 neurons to distal CA1 theta oscillations selectively predicts memory performance

**DOI:** 10.1101/2023.06.21.546025

**Authors:** Shih-Pi Ku, Erika Atucha, Nico Alavi, Motoharu Yoshida, Joszef Csicsvari, Magdalena M. Sauvage

**Affiliations:** Leibniz Institute for Neurobiology, Functional Architecture of Memory Dept., Magdeburg, Germany; Institute of Science and Technology (IST), Klosterneuburg, Austria; German Center for Neurodegenerative Diseases (DZNE), Magdeburg, Germany; Otto von Guericke University, Medical Faculty, Functional Neuroplasticity Dept., Magdeburg Germany; Center for Behavioral Brain Sciences (CBBS), Magdeburg, Germany

**Keywords:** Recognition memory, proximal, distal, hippocampus, CA1, CA3, theta, support vector machine, population coding, odor

## Abstract

How the coordination of neuronal spiking activity and brain rhythms between hippocampal subregions supports memory function remains elusive. We studied interregional coordination of CA3 neuronal spiking activity with CA1 theta oscillations by recording electrophysiological signals along the proximodistal axis of the hippocampus in rats performing a high memory demand recognition memory task adapted from humans. We found that CA3 population spiking activity occurs preferentially at the peak of distal CA1 theta oscillations only when animals recalled previously encountered stimuli. In addition, decoding analyses revealed that only population cell firing of proximal CA3 together with that of distal CA1 can predict memory performance in the present non-spatial task. Overall, our work demonstrates an important role of the synchronization of CA3 neuronal activity with CA1 theta oscillations for successful recognition memory.

## Introduction

Retrieving memories for previously encountered objects, odors, people or places is crucial for dealing with daily-life situations. In humans and animals, this ability relies heavily on the hippocampus, which is located in the medial temporal lobe of the brain in humans and rodents (Eichenbaum et al., 2007, Koen and Yonelinas, 2014, Yonelinas et al., 2010, Scoville and Milner, 1957, Zola-Morgan et al., 1982, Kaada et al., 1961). The hippocampus comprises distinct functional domains, such as the hippocampal subfields CA1 and CA3 that are part of the anatomical pathway the most scrutinized for its contribution to memory (the ‘trisynaptic loop’) which features heavy anatomical projections from CA3 to CA1 (Amaral and Witter, 1989, Amaral, 1993, Ishizuka et al., 1990, Brivanlou et al., 2004, van Strien et al., 2009). An important focus in memory research has been to elucidate the mechanisms supporting memory. How the neuronal spiking activity coordinates local and between brain areas’ rhythms has drawn much attention (Zutshi et al., 2022, Fernandez-Ruiz et al., 2017, Schomburg et al., 2014, Montgomery et al., 2008, Dragoi et al., 1999, Colgin et al., 2009, Zheng et al., 2016, El-Gaby et al., 2021). However, whether these functional relationships contribute to memory performance in a decisive manner remains unclear.

Theta oscillation (centered at 8Hz) is prominent in the hippocampus and plays an important role in memory function (Buzsaki and Moser, 2013, Buzsaki, 2005). In rodents, theta oscillations facilitate memory processes by coordinating signals within or between areas as seen with the precession of neuronal firing in relation to theta phase as a function of animals’ experience (Skaggs et al., 1996, Dragoi and Buzsaki, 2006, Colgin, 2013, Burgess and O’Keefe, 2011, Harris et al., 2002) or illustrated by the synchronization of spiking activity with theta oscillations between the hippocampus and the prefrontal or the entorhinal cortex (Benchenane et al., 2010, Hyman et al., 2010, Ishino et al., 2017, Siapas et al., 2005, Wirt and Hyman, 2019 and (Mizuseki et al., 2009, Frank et al., 2001, respectively). In humans, the amplitude of theta oscillations varies in a task-dependent manner and memory performance improves when stimuli are presented at a theta frequency (Pacheco Estefan et al., 2021, Leszczynski et al., 2015, Axmacher et al., 2010, Canolty et al., 2006).

A tremendous amount of knowledge has been accumulated on local couplings within the hippocampal subfields CA1 or CA3 (i.e. spike-to-theta phase locking within CA1 or CA3) in rodents in studies investigating spatial navigation using tasks such as running on a linear track (Buzsaki and Eidelberg, 1983, Fox et al., 1986, Harris et al., 2003, Buzsaki and Tingley, 2018, Dragoi and Buzsaki, 2006, Harris et al., 2002). In contrast, the relationship between theta oscillations and neural spiking activity between hippocampal subregions and its ties to memory performance are not yet well-understood. Hence, whether the coordination of CA3 neuronal spiking activity with CA1 theta oscillations (i.e. CA3 spike to CA1 theta phase coupling) bears an important role in recognition memory remains elusive, especially in tasks that are devoid of salient spatial information with high memory load which are often used in humans (Radvansky, 2006, Yonelinas, 1997, Wixted et al., 2018, Stark and Squire, 2000).

Importantly, immediate-early gene imaging and electrophysiological evidence for a functional segregation along the proximodistal axis of CA1 and CA3 in terms of spatial versus object/odor information processing has accumulated during the past decade (Beer et al., 2018, Flasbeck et al., 2018, Nakamura et al., 2013, Henriksen et al., 2010, Lu et al., 2015, Burke et al., 2011, Vandrey et al., 2021). These findings suggest the existence of distinct subnetworks within the hippocampus that would preferentially process spatial or non-spatial information: the ‘spatial’ subnetwork including distal CA3 (distCA3) and proximal CA1(proxCA1) and the ‘non-spatial’ subnetwork including proxCA3 and distCA1 (Fig 1A; Nakamura et al., 2013; Beer et al., 2018). However, *in-vivo* evidence for such subnetworks as well as evidence of a functional interaction within these subnetworks are missing.

**Figure 1.**
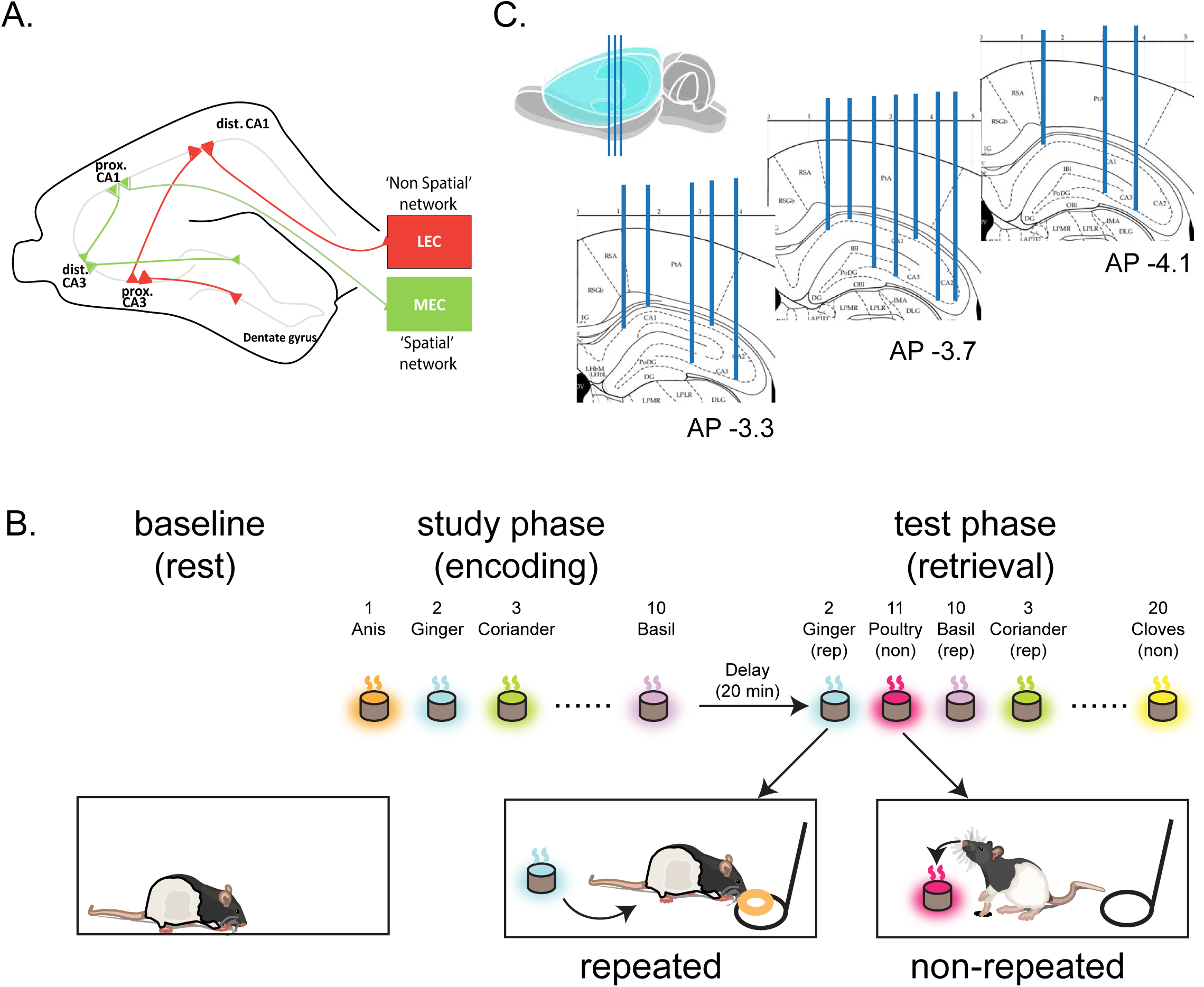
Schema of the ‘spatial’ and ‘non-spatial’ Medial Temporal Lobe subnetworks, odor recognition memory paradigm and hippocampal recording sites. A) The ‘segregated paths’ model describing subnetworks preferentially processing spatial and non-spatial information: the ‘spatial’ and the ‘non-spatial’ subnetworks (in green and red, respectively; Nakamura et al., 2013, Beer et al., 2018). The ‘non-spatial’ subnetwork includes the lateral entorhinal cortex (LEC), proximal CA3 (proxCA3), distal CA1 (distCA1) and the exposed blade of the dentate gyrus (exp. DG). The ‘spatial’ subnetwork includes the medial entorhinal cortex (MEC), distal CA3 (distCA3), proximal CA1 (proxCA1) and the enclosed blade of the DG (encl. DG). B) Delayed non-matching to odor memory task: each day, the session starts with rats resting in their home cage placed in the recording booth for 20 minutes (baseline). Thereafter, rats sampled 10 different odors, one by one, during the study phase (the study list changes each day). After a 20-minute delay, the memory for these odors (‘repeated’ odors) is tested by presenting to the rats the same odors intermixed with 10 other odors that were not experienced during the study phase (i.e. ‘non-repeated’ odors), one by one and in a pseudorandom manner. Both ‘repeated’ and ‘non-repeated’ odors are part of a pool of 40 familiar odors that are used in a pseudorandom manner over the 2 months of training. According to a non-matching to sample rule, rats report a ‘repeated’ odor by refraining digging and turning away from the stimulus cup (i.e. a correct response) and a ‘non-repeated’ odor by digging into the stimulus cup (i.e. a correct response). Memory performance (% correct) is calculated on the basis of these responses. C) Recording sites are located at the tip of the blue parallel lines and distributed along the entire proximodistal axis of CA1 and CA3 at about AP-3 to -4 from Bregma.

Here, we studied *in-vivo* the relationship between neuronal activity and theta oscillations within and between CA3 and CA1 and its ties to memory performance. In an attempt at bridging further human and animal recognition memory, we leveraged a high demand memory task adapted from humans to rats (Fortin et al., 2004, Nakamura et al., 2013, Sauvage et al., 2010), simultaneously recorded the electrophysiological activity along the proximodistal axis of CA1 and CA3 and analyzed neuronal activities at population and single cell levels. In humans, recognition memory tasks typically consiste in presenting a list of stimuli, for example words appearing one at a time in the center of a screen, consecutively to which memory for these stimuli is evaluated. This is done by presenting the same stimuli, one at a time, intermixed with stimuli that have not been presented on the same day and by calculating the percentage of correct responses (Radvansky, 2006, Yonelinas, 1997, Wixted et al., 2018, Stark and Squire, 2000). Likewise, here, the memory for a stimulus list presented during a study phase was tested in rats by presenting at test the same stimuli (‘repeated’ stimuli) intermixed with stimuli that rats had not experienced during the study phase (‘non-repeated’ stimuli) (Fig. 1B). Given that olfaction is the primary sense of rodents and that rats have an exceptional memory capacity for odors, odors from a pool of fourty common household scents were used as stimuli.

We report that CA3 neurons were selectively phase-locked close to the peak of distal CA1 theta oscillations during the retrieval of the memory of a previously encountered stimulus (‘repeated’ stimulus) but not when the stimulus was not experienced during the study phase (‘non-repeated’ stimulus). We also show that the population phase information of CA3 spikes predicts memory performance trial-by-trial only when theta was referenced to the distal part of CA1. Our results suggest that the coordination of CA3 activity with CA1 theta rhythm might constitute one of the fundamental mechanisms supporting memory retrieval and indicate a preferential involvement of proximal CA3 and distal CA1 in processing non-spatial memory.

## Results

Local field potential (LFP) and single-unit activities in CA1 and CA3 were simultaneously recorded in the same animals (5 rats; 633 neurons, in total 394 in CA1 and 239 in CA3, respectively; Table 1) from up to 16 sites using tetrode arrays covering the transverse (proximodistal) axis of CA1 and CA3 (Fig 1C). The localization of the tetrodes and clustering criteria are reported in the methods section. Population and single cell firing, characteristics of theta oscillations (amplitudes) as well as cross (CA3-CA1) or within area (CA1-CA1 or CA3-CA3) spike-theta phase-locking properties and their relationships to memory performance were studied in the same behavioral arena: at baseline (i.e. prior to the study phase) as well as during the study and the test phase of the task (Fig. 1B; analysis epochs are described in Methods and Supp. Fig 1).

**Table 1.**
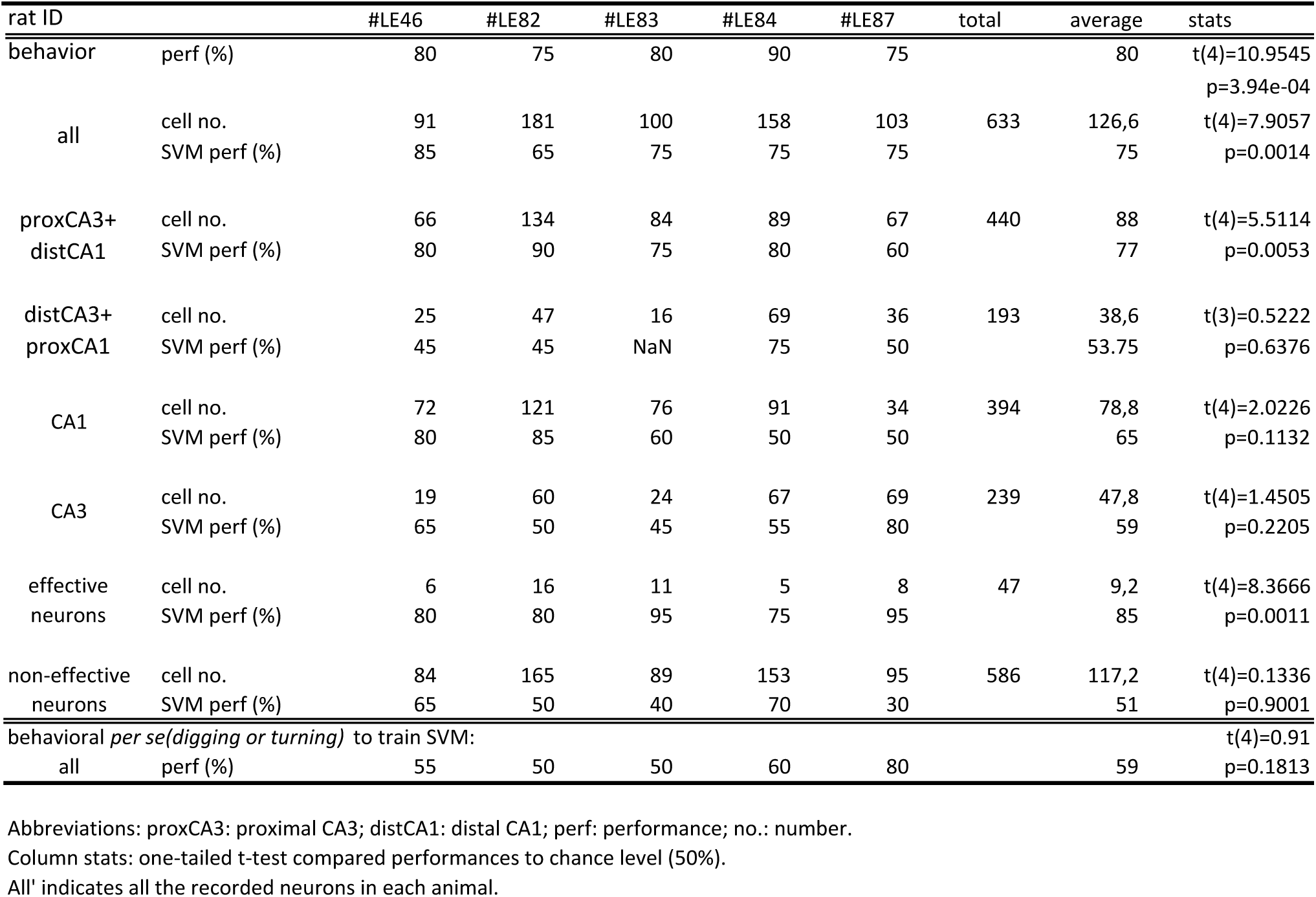
Memory performance, population classification performances and number of single cells analyzed for each animal

### A proximodistal gradient in population cell firing activities discriminating ‘repeated’ from ‘non-repeated’ odors trial by trial

Discrimination between stimuli or spatial contexts has been recently reported to be coded at the population level rather than by individual cell firing (El-Gaby et al., 2021, Guzowski et al., 2004, Leutgeb et al., 2004). Hence, we first investigated if population firing activities, i.e. the combined mean firing rate of individual neurons, could reflect discrimination between ‘repeated’ and ‘non-repeated’ odors during memory retrieval, trial-by-trial. We used Support Vector Machine (SVM) to analyze the population firing activities of all recorded neurons in each animal (n=127 neurons in average per animal; Table 1; see Methods for details; (Ku et al., 2008, Brinkmann et al., 2015). The SVM classification performance of all neurons matched animals’ memory performance, which is in agreement with a contribution of hippocampal population cell firing activities to memory retrieval (averaged SVM performance all neurons: 75% correct; averaged animal memory performance: 80% correct, n=5; SVM vs memory performance: t(8) = 1.1952, p = 0.2662, 2-sample t-test; Table 1; SVM > chance level: *t*(4) = 7.9057, *p* = .0014, one-tailed t-test, Bonferroni corrected; Fig. 2A-B, dark blue and white bars, respectively). In addition, we observed that only population cell firing of the areas belonging to the hippocampal subnetwork preferentially processing non-spatial information (proxCA3 + distCA1) could account for the classification performance of all recorded neurons, suggesting a preponderant role of cell firing at the population level in proxCA3 and distCA1 for odor memory retrieval (SVM performance (proxCA3+distal CA1): 77%; compared to all neurons: t-test, *t*(8)= 0.343, *p* = 0.7407; > chance level: *t*(4) = 5.5114, p = .0053, one-tailed t-test, Bonferroni corrected, proxCA3+ distCA1 neurons represents ∼69% of the recorded neurons (440/663), see Table 1; Fig. 2B, red bar, and Methods for details). This result was further supported by the finding that any other combination of areas studied yielded a chance level SVM performance. For example, when only neurons belonging to the subnetwork preferentially processing spatial information were considered (distCA3 + proxCA1, representing 31% of the recorded population (193/633, table 1) or when only CA1 or CA3 neurons were considered (distal + proximal parts; 62 % (394/633) and 38% (239/633) of the recorded population, respectively; SVM performance: distCA3 + proxCA1: 53.75%; *t*(3)= 0.5222, *p* = 0.6376; CA1 = 65%, *t*(4) = 2.0226, *p* = 0.1132; CA3: 59%, *t*(4) = 1.4505, *p* = 0.2205; all one-tailed t-tests to chance level, Fig. 2B, grey bars; Table 1). Further analysis see Supp. Results, Supp. Fig. 2A). Moreover, the SVM classifier for the areas of the counterpart subnetwork (distal CA3+proximal CA1) performed significantly lower than the classifier for proxCA3+distCA1 (2-sample one-tailed t-test, t(7)=2.7664, p=0.0139), supporting the hypothesis that proxCA3 and distCA1might be more engaged than distCA3 and proxCA1 for retrieving non-spatial (odor) memory. Furthermore, we tested whether population firing activities of all neurons could predict behavioral responses *per-se* (i.e. digging or turning independently of whether animals responded correctly or not). As a reminder, the period analyzed started upon stimulus presentation (when the front edge of the stimulus platform crossed the wall of the cage) and ended 2sec later before behavioral response (turning or digging; the time-window of 2 sec was defined based on prior experiments showing that behavioral response in this task occurred shortly after 2 sec, Supp. Fig. 1). Rats were sniffing and alertly waiting during these 2-sec periods. We found that population firing activities of all neurons did not account for digging or turning *per-se* (averaged SVM performance: 59%, see Table 1, n=5, SVM > chance level: t(4)=0.91, p = 0,1813).

**Figure 2.**
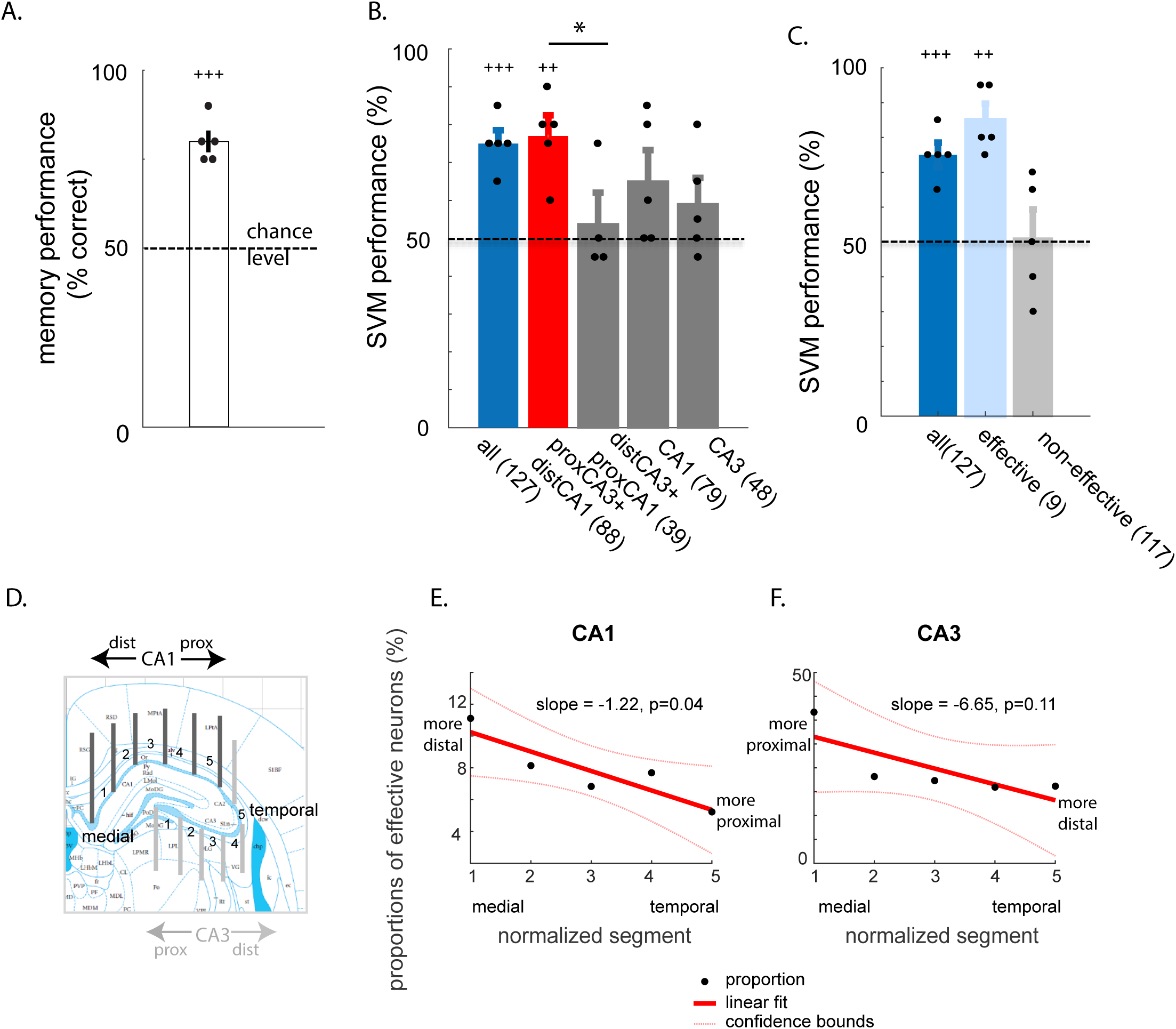
Memory performance, population classification performance of neurons discriminating between ‘repeated’ and ‘non-repeated’ stimuli and topographical distribution of ‘effective’ neurons along the proximodistal axis of CA1 and CA3. A) Memory performance on the recording day: rats were trained until they reached a plateau performance of at least 75% correct choice over 3 consecutive days, subsequently to which single cell firing and LFP were recorded during an additional session with above-threshold (75% correct) behavioral performance. Dots indicate individual performance (N=5). B) Population classification performances across the 5 recorded rats obtained using Support Vector Machine (SVM) based on either: all neurons, neurons belonging to the ‘non-spatial’ subnetwork (proximal CA3+distal CA1), to the ‘spatial’ subnetwork (distal CA3+proximal CA1) or to CA1 or CA3 (both proximal+ distal parts). Only the population classification performance of the neurons belonging to the ‘non-spatial’ subnetwork (proxCA3+dist CA1) is comparable to memory performance and the SVM performance of all recorded neurons. Moreover, SVM performance of proxCA3+dist CA1 is higher than that of neurons belonging to the ‘spatial’ subnetwork (distCA3+proxCA1). Averaged cell counts per animal are in brackets and rounded up or down for clarity purpose (see Table 1). C) The SVM performance of ‘effective’ neurons (in average n=9 neurons per rat) can also account for the SVM performance of all recorded neurons and memory performance, not that of the ‘non-effective’ neuron population. D) Anatomical location of the five proximodistal segments of CA1 and CA3 in which neural activities were recorded. Segments 1 are more medial, segments 5 more temporal (see Methods for segment normalization procedure). In CA1, segment 1 is the most distal and segment 5 the most proximal; in CA3, segment 1 is the most proximal, segment 5 the most distal. E and F) Distribution of the proportions of ‘effective’ neurons along the proximodistal axis of either CA1 (E) or CA3 (F). In CA1, the slope of the linear regression is significantly smaller than zero (p=0.04), indicating a distal to proximal gradient of the distribution of effective neurons in CA1 (higher proportion in distal CA1). In CA3, although the slope failed to reach significance (p=0.11), comparing the two most proximal (1,2) to the two most distal (4,5) segments of CA3 indicated a larger proportion of ‘effective’ neurons in the most proximal part. Altogether, these data indicate a preponderant role of population cell firing in proximal CA3 and distal CA1 (the ‘non-spatial’ subnetwork) during odor memory retrieval when compared to that of distal CA3 and proximal CA1, parts of the ‘spatial’ subnetwork. dots: proportions of effective neurons in each segment; red line: linear fit; red dotted lines: 95% confidence bounds. Error bars: S.E.M. *: p<0.05; ++: p<0.01;+++:p<0.001.

We also showed that all recorded neurons did not contribute to the same extent to the population classification performance by using an iterative feature selection procedure, deleting neurons one at a time to assess if the deletion of a specific neuron affects the population performance (Supp. Fig. 3) For each iteration, one neuron was removed from the whole population (n=N) in a given animal and the rest of neurons (n=N-1) underwent SVM classification analysis. If the performance of the rest of the neurons (n=N-1) was lower than the performance while taking all the neurons (n=N) into account, the neuron was termed ’effective’ neuron, if not: ‘non-effective’ neuron (see Supp. Fig. 3 for the iterative procedure and Methods for details). In average 7% of the neuronal population recorded in each animal were ‘effective’ neurons (i.e. in average 9 effective vs 117 non-effective neurons per animal, 47 vs 586 in total across animals; Table 1). The classification performance across animals for these effective neurons was comparable to that of all recorded neurons (mean SVM performance effective neurons= 85% correct; : *t*(8) = 1.9069, *p* = 0.093, 2-sample t-test; one-tailed t-test to chance level: *t*(4) = 8.3666, *p* = .0011; Table 1, Fig. 2C light blue bar). In contrast, SVM performance of the remaining non-effective neurons did not differ from chance level (mean SVM performance non-effective neurons=50.1% correct; one-tailed t-test to chance level: t(4) = 0.1336, p = 0.9001; Table 1, Fig. 2C, light grey bar). These results indicate that population firing activities of about 7 % of the recorded neurons (47/633, effective neurons/all neurons across animals) can account for the SVM classification performance of all recorded neurons (Fig. 2C, Table 1). Moreover, the topographical distribution of the effective neurons was not homogenous along the proximodistal axis of CA1 and CA3 (Supp. table2). In CA1, the distribution of effective neurons linearly decreased from distal to proximal (Fig. 2D-E; p=0.04, slope different from zero; Supp. Table 2). In CA3, although the distribution did not linearly decrease from proximal to distal when all segments were taken into consideration (p=0.11), a direct comparison between the two most medial and the two most temporal segments revealed a higher proportion of effective neurons in the proximal than the distal part of CA3 (Fig 2F; segments 1-2 vs segments 4-5: p= 0.0012, z= 3.2297, two-proportion z-test), bringing further support to the hypothesis of a preferential tuning of proxCA3 and distCA1 (belonging to the ‘non-spatial’ subnetwork) to the processing of odor information during memory retrieval.

Finally, we examined whether there was a general increase or decrease in firing rates for the retrieval of ‘repeated’ odors when compared to ‘non-repeated’ odors in individual CA1 and CA3 neurons. Mean individual firing rates differed between ‘repeated’ and ‘non-repeated’ odors in only 41 out of 633 neurons (i.e. 6.5% of all neurons, 8 neurons per animal in average, Supp. Table 3; paired t-test, threshold at p< 0.05, example cell firing patterns in Supp. Fig. 4,. see Methods for details; 5 out of the 41 are effective neurons) and did not generally increase or decrease for either stimulus type (4.8 neurons per animal increased and 3.4 decreased, increasing vs decreasing proportion: p=0.27, two-proportion z-test, see Supp. Table 3 for individual animal). Combining the firing rates of the neurons with differential firing rates could predict memory performance (SVM performance: 61%, t(4)=11, p=1.94×10^-04^, one-tailed t-test to chance level; Supp. table1) but to a lower extent than combining the firing rates of effective neurons (comparable in number: n=9 per animal in average, 47 in total across animals; comparison to SVM performance of effective neurons (i.e. to 85% correct); t(8)=5.5799, p=5.22×10^-4^, 2-sample t-test). Furthermore, we assessed whether the firing rate, this time, of all CA1 and CA3 neurons (i.e. differentially firing or not) were overall enhanced or suppressed during repeated trials in comparison to non-repeated trials. This was not the case as CA1 neuronal firing rates were higher for the retrieval of ‘repeated’ odors than ‘non-repeated’ odors in only two out of five rats (no differences in CA3 in these rats; no differences in the three remaining rats in either CA1 or in CA3; Supp. Fig. 5). This together with our earlier finding that the combination of population firing activities in proxCA3+distCA1 can discriminate repeated from non-repeated odors (see SVM analyses) indicate that differences in population firing activities between stimulus types might be a better predictor of memory performance than a general enhancement or suppression of firing rates.

Altogether, these results show that population cell firing activities in distCA1 and proxCA3 alone can account for the population classification performance of all recorded neurons, suggesting that the ‘non-spatial’ hippocampal subnetwork might play a preponderant role in memory retrieval in this task. Moreover, this finding raises the possibility of a preferential coordination of proxCA3 and distCA1 electrophysiological signals during odor memory retrieval. To further test this hypothesis, we investigated CA3 spikes to CA1 theta phase-locking along the proximodistal axis of CA1 and CA3.

### Preferential CA3 spike-distal CA1 theta coordination during odor memory retrieval

Theta oscillations in CA1 have been shown to play a crucial role in rodents in spatial navigation tasks while, in comparison, much less is known about their role in non-spatial memory, especially in tasks with high memory demands. Hence, we first studied CA1 theta oscillation characteristics (cycle-by-cycle amplitudes, i.e. power) in the odor task used in the present study. This was done along the proximodistal axis of the hippocampus in the animals previously described. Second, we investigated the relationship between neuronal spiking and theta oscillations between and within CA3 and CA1 and how this relationship could account for memory performance.

#### a) CA1 theta oscillations are strong during odor memory, especially during retrieval

Theta oscillations in distCA1 were strong during all phases of the task: i.e. at baseline (resting period before study), study and retrieval (see Fig. 3A-B for individual examples of theta oscillations and power spectra and Supp. Fig. 6 for examples of longer raw traces). At the group level, the frequency of theta oscillations were similar throughout the phases of the task (F(2,12)=0.07, p=0.9303, one-way anova, Supp. Table 4). In addition, using a cycle-by-cycle analysis optimized for trials with short durations, as is the case in our task (∼2sec; (Cole and Voytek, 2019)), we showed that averaged theta cycle amplitudes (i.e. theta power) were significantly larger at retrieval than study or baseline in all animals in CA1 (all ps<0.0107, Mann-Witney U-test due to the non-nomal distribution of theta amplitudes, Bonferroni corrected; see an example of theta cycle amplitudes in Fig 3C; no significant differences between baseline and study phase, all p>0.02; Bonferroni corrected; Supp. Table 5). In contrast, theta power was comparable between retrieving ‘repeated’ and ‘non-repeated’ odors in 4 out of 5 animals (p>0.068, Mann-Witney U-test, Fig.3D, see Supp. Table 5 for individual animals) and between ‘pre’- and ‘in-trial’ periods of the task in 3 out of 5 rats (i.e. 2 sec before vs. after stimulus delivery at retrieval, respectively: p > 0.05; Supp. Table 5, see Supp. Fig. 1 for the description of pre-and in-trial period and Methods for details) suggesting that theta power is rather homogeneous throughout the retrieval phase of the task.

**Figure 3.**
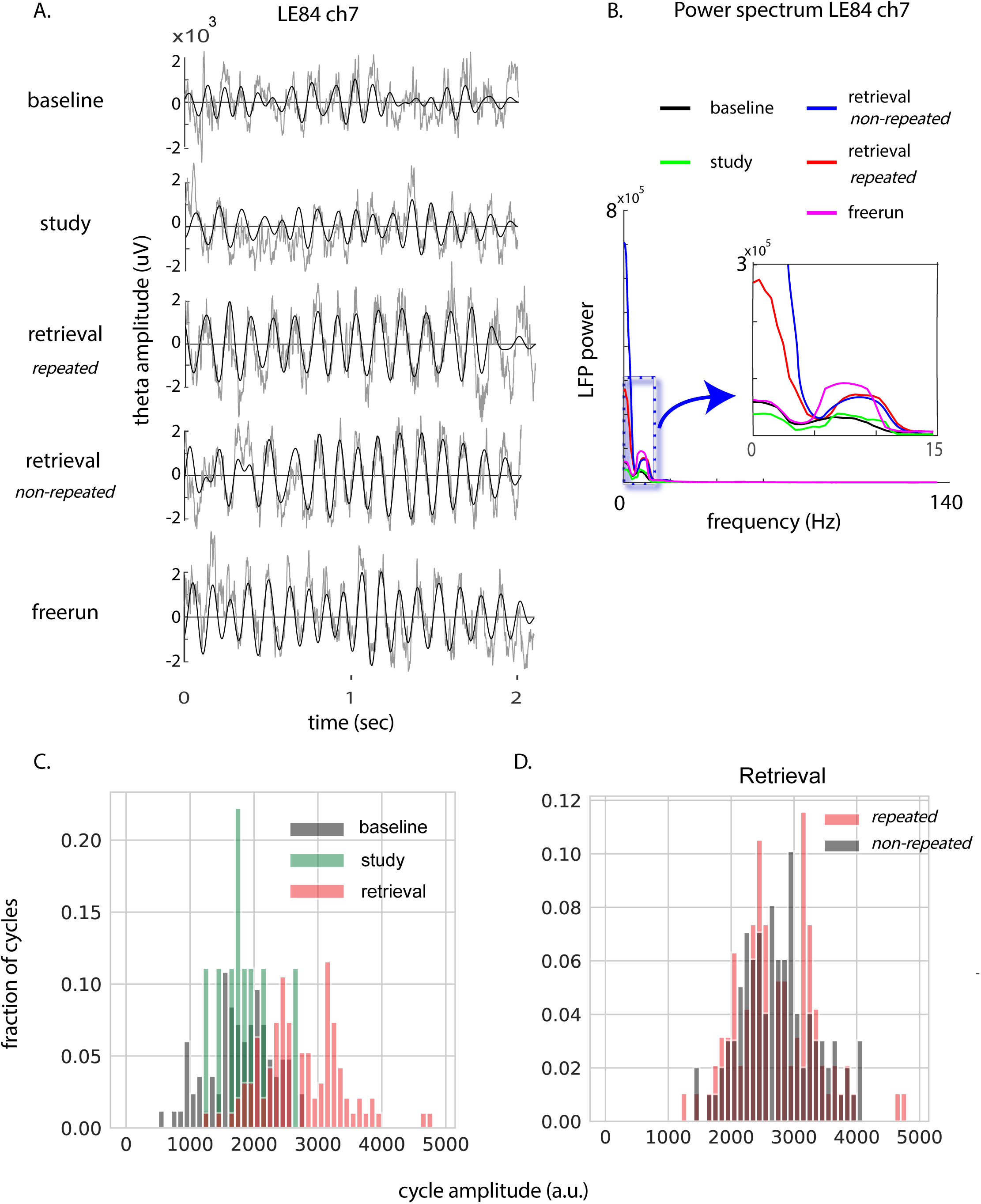
Theta power is high during non-spatial (odor) memory, especially during retrieval. A) Examples of individual unfiltered local field potential overlaid with filtered theta oscillations (6-12Hz) of one representative recording session during baseline, study, retrieval and freerun, indicating that theta oscillations are strong during the different phases of the present task that is devoid of salient spatial components (averaged animal’s speed at baseline: 1.63cm/sec; speeds for retrieval and freerun traces are matched: ∼5.5 cm/sec; animal’s speed at study: 8,48 cm/sec. Note that even when animal’s speed is higher (study), theta amplitudes are lower. B) Power spectrum analysis of one example session indicating a higher theta power at retrieval (for both ‘repeated’(red) and ‘non-repeated’(blue) trials) and freerun (magenta) than during baseline (black) and study (green). C) Example distributions of the amplitude of each theta cycle during baseline, study and retrieval in one representative animal. Cycle-by-cycle analyses optimized for short duration trials (Cole and Voytek, 2019) revealed that averaged theta cycle amplitudes (i.e. theta power) were enhanced during retrieval compared to baseline or study. D) In contrast, theta power was comparable during the retrieval of either type of stimulus as revealed by the overlapping distribution of theta cycles’ amplitudes for ‘repeated’ and ‘non-repeated’ odors. A.u.: arbitrary unit.

Because running speed was reported to affect theta power, we further studied the relationship between the speed of animals in our task and the amplitude of theta oscillations. The average speed significantly differed across task phases (F(2,12)=30.12, p=2.1×10^-5^, one-way ANOVA, Supp. Table 6). Post-hoc analyses revealed that the speed at study and test was very low and comparable across the five animals (median+-std=4.34+-0.03 and 4.69+-0.26cm/sec, respectively; 2-sample t-test, t(8)=1.4118, p=0.2595) and, as expected, higher than at baseline (1.35+-1.38cm/sec; baseline vs study or test: p< 4.97×10^-04^, 2-sample t-tests: Bonferroni corrected, see Supp. Fig7 and Supp. Table 6 for individual data). Moreover, regression analyses showed overall the lack of a significant relationship between speed and theta oscillation amplitudes during retrieval and study (all ps>0.119, but for 2 out of the 5 rats at study; both ps< 0.017; see Supp. Fig.8 and supplementary results for more details), suggesting that speed alone does not account for the enhancement of theta oscillations’ amplitudes observed between retrieval and baseline or study.

Altogether, these results indicate that theta power is high during the present odor memory task, as it is the case in spatial tasks. Moreover, our data show that theta power in distal CA1 is especially high during memory retrieval, that this enhancement lasts throughout the entire retrieval phase of the task and that the increase in theta oscillations’ amplitude is unlikely to stem solely from changes in the speed of the animals.

#### b) Theta power is higher in distal CA1 than proximal CA1 in trained, but not naïve, rats

Analysis of theta oscillations’ amplitudes in the five trained rats revealed higher theta amplitudes in distal than proximal CA1 independently of the phase of the task (Friedman’s test on Kolmogorov–Smirnov (KS) test D-values (difference of theta amplitudes between dist. and prox.CA1): p=0.2466; see Fig. 4A-C for examples of theta amplitudes’ distribution, Supp. Table 8, see Methods for details). Such a proximodistal difference was also observed during freerun in trained rats (all ps<2.43×10^-23^, KS test, Bonferroni corrected, Fig. 4D, Supp. Table 7&8; see Supp. results for more analyses on freerun). In a striking contrast, amplitudes of theta cycles did not consistently differ between distal and proximal CA1 in naive animals resting in their cage (n=5; Fig. 4E, Supp. Table 7&8) and a direct comparison between trained and naive animals at baseline established that proximodistal differences were significantly larger in trained than naive animals (p=0.0079, Wilcoxon signed-rank test on KS test D-values, Fig. 4F, Supp. Table 8). Importantly, the average speed of the animals at baseline was comparable between trained and naïve rats (1.35+-1.18 and 0.99+-1.08cm/sec, respectively; 2-sample t-test, p=0.6258, Supp. table6), indicating that the increase in theta amplitudes in trained rats unlikely stems from a difference in animals’speed.

**Figure 4.**
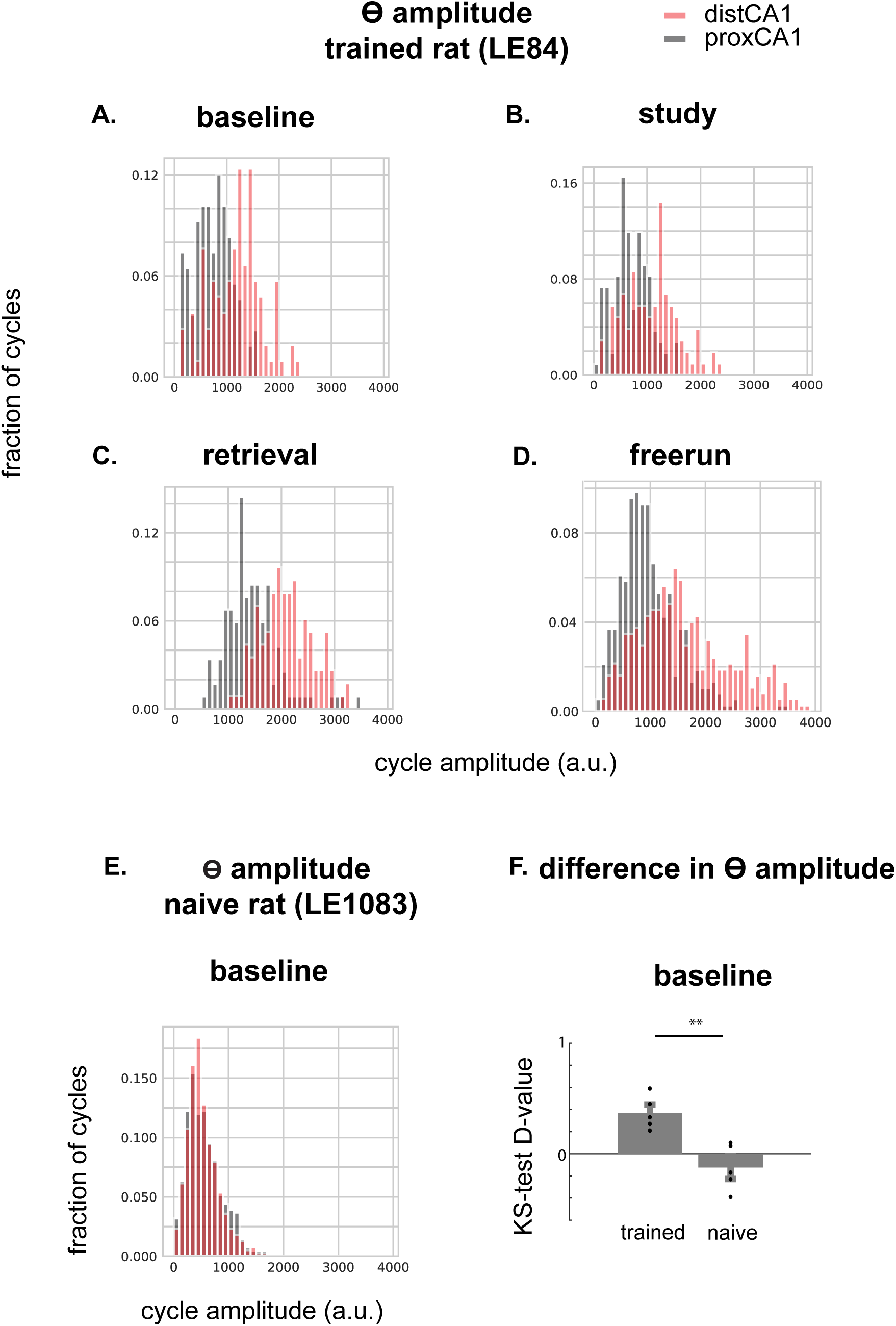
The distributions of theta cycles amplitudes in distal and proximal CA1 in representative trained or naïve rats. A-D) the amplitudes of theta cycles (power) are enhanced in distal CA1 when compared to proximal CA1 in all phases of the task as well as during freerun when the animal has been trained (over 2 months) on the odor memory task. E.) In contrast, theta power is comparable in distal and proximal CA1 in naïve animals during baseline (resting period). F.) Differences between distal and proximal CA1 in theta cycle amplitudes are obtained using the Kolmogorov–Smirnov test to generate the D value (Maximum|Fo(X)−Fr(X)|, F(X)=observed cumulative frequency distribution of a random sample of n observations) for each animal. Theta cycles’ amplitudes at baseline are higher in distal than proximal CA1 in trained (N=5) but not in naïve animals (N=5). Of note, the average speed of trained and naïve rats at baseline is comparable (**: p<0.01; dots: individual D-values; error bars: S.E.M. Altogether, these results suggest the absence of ‘constitutive’ proximodistal differences in theta power along CA1 and might indicate a tuning of distal CA1 to the processing of non-spatial (odor) information over time.

Altogether these results suggest a selective enhancement of theta oscillations in distal compared to proximal CA1 in terms of amplitudes of theta cycles in rats that underwent over 2 months of training in the odor memory task used in the present study. In addition, these data reveal the absence of gross ’constitutive’ differences between distal and proximal CA1’s theta oscillations suggesting that theta amplitudes enhancement in distal CA1 might stem from a selective tuning of this area to the processing of odor across and within sessions.

#### c) Only CA3 spikes’ phase-locking to distal CA1 theta can account for memory performance

To test whether proximodistal differences in the coordination of neuronal spiking and theta oscillations between CA3 and CA1 play an important role in memory retrieval in the present task, we evaluated the proportion of CA3 neurons phase-locked to theta oscillations in either distal or proximal CA1. Single cell level analyses revealed that the vast majority of the CA3 neurons that spiked during the retrieval of both ‘repeated’ and ‘non-repeated’ odors are phase-locked to distal CA1 theta (135/150; i.e. 90%, 27 per animal in average). For about half of these neurons (69/135, i.e. 51%, out of which 10 effective neurons, 13.8 per animal; Fig. 5A left), the preferred phase-locking angle differed between ‘repeated’ and ‘non-repeated’ odors (threshold at p <0.05, circular Watson-Williams multi-sample test for equal variance, see methods for details). These neurons were termed ‘phase discriminating’ neurons Most phase discriminating neurons (i.e.71%; 49/69; 9.8 neurons per animal, Fig. 5A right) were locked to different phases of distal CA1 theta when retrieving ‘repeated’ or ‘non-repeated’ odors (see individual example in Fig. 5B,C) while the remaining were phase-locked to only one of the two stimulus type (‘non-repeated’ odors: 19%;13/69 ; 2.6 neurons per animal; ‘repeated’ odors: 10%;7/69 ; 1,4 neuron per animal; Fig. 5D,E and F,G, respectively). Similar results were found when CA3 spikes phase-locking to proximal CA1 theta was studied (89% are phase-locked;136/150 ; 27.2 per animal) among which 46% (62/136 ; 12.4 per animal) are discriminating neurons. In summary, single cell level analyses showed that the spikes of a similar proportion of CA3 neurons were phase-locked to distal or proximal CA1 theta.

**Figure 5.**
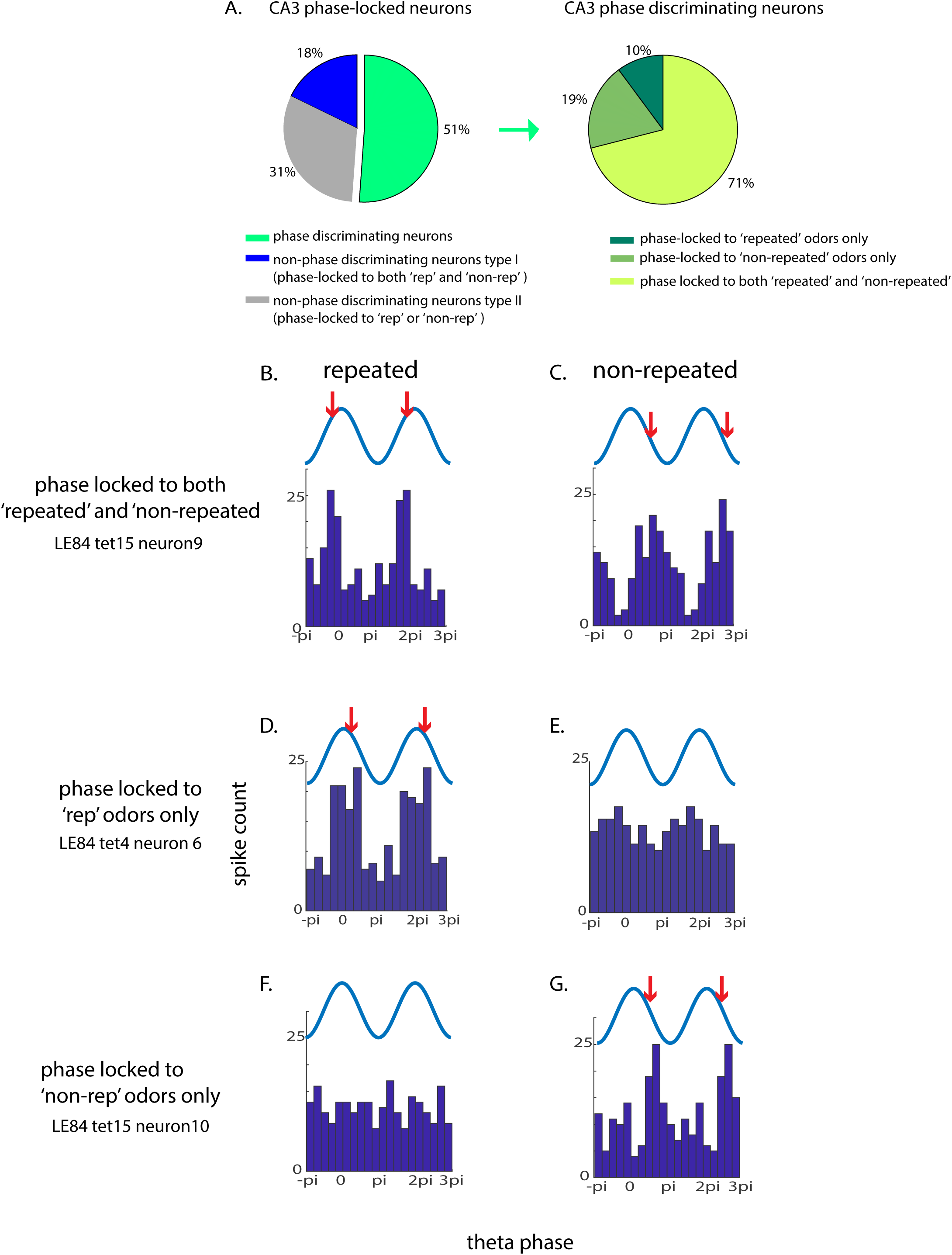
The majority of CA3 spikes are phase-locked to distal CA1 theta oscillations during memory retrieval. A) Left: proportion of different types of CA3 neurons phase-locked to distal CA1 theta oscillations: the ‘phase discriminating neurons’ (in green, with a phase-locking angle differing between ‘repeated’ and ‘non-repeated’ odors) and non-phase discriminating neurons: type I (in blue) and type II (in grey). Type I neurons are phase-locked to the same angle of the theta phase oscillations for ‘repeated’ and ‘non-repeated’ odors. Type II are phase-locked for only one type of stimulus (grey). About half of the CA3 neurons phase-locked to CA1 theta (51%) can discriminate between ‘repeated’ and ‘non-repeated’ stimuli, i.e are phase-discriminating neurons. Right: CA3 phase-discriminating neurons (green on the left) can be subdivided into three groups: the majority of neurons (>70%) are phase-locked to different phases of distCA1 theta oscillations for ‘repeated’ and ‘non-repeated’ odors while about 20% are phase-locked for ‘non-repeated’ and 10% for ‘repeated’ stimuli. B-C) Phase distributions of representative neurons phase-locked to CA1 theta with different angles for the retrieval of ‘repeated’ and ‘non-repeated’ odors; D,E) phase-locked only for the retrieval of ‘repeated’ odors; F,G) phase-locked only for the retrieval of ‘non-repeated’ odors (F,G). Red arrows: mean angle of the population cell firing. ‘rep’: repeated; ‘non-rep’: non-repeated.

At the population level, the analysis of the mean angle of the phase-discriminating CA3 neurons to distal CA1 theta revealed that CA3 neurons tend to fire slightly after the peak of distal CA1 theta (circular mean = 1.5°) only for the retrieval of ‘repeated’ odors (circular Rayleigh test: ‘repeated’: *p* =.0168; ‘non-repeated’: *p* = .88; Fig. 6A-B). In contrast, using theta oscillations in proximal CA1 as a reference did not yield any specific patterns for ‘repeated’ or ‘non-repeated’ odor retrieval (circular Rayleigh test: repeated: *p* = 0.7; non-repeated: *p* =0.56; Fig. 6C-D). To directly evaluate whether CA3 spikes were differentially phase-locked to distal or proximal CA1 theta for either ‘repeated’ or ‘non-repeated’ odors, we compared the variance of the distributions of the mean theta phases of CA3 spikes between distal and proximal CA1 for repeated or non-repeated odors separately. The variance of the mean phases of CA3 spikes using distal CA1 as a reference was lower than the variance obtained using proximal CA1 theta. This was the case only for ‘repeated’ odor trials (Watson-Williams multi-sample test for equal variance; ‘repeated’: F(1,150)=13.49, p= 0.0003; ’non-repeated’: F(1,150)=3.42, p=0.0665), further suggesting a selective role of the phase-locking of CA3 neurons to distal CA1 theta for retrieving the memory of non-spatial (odor) information.

**Figure 6:**
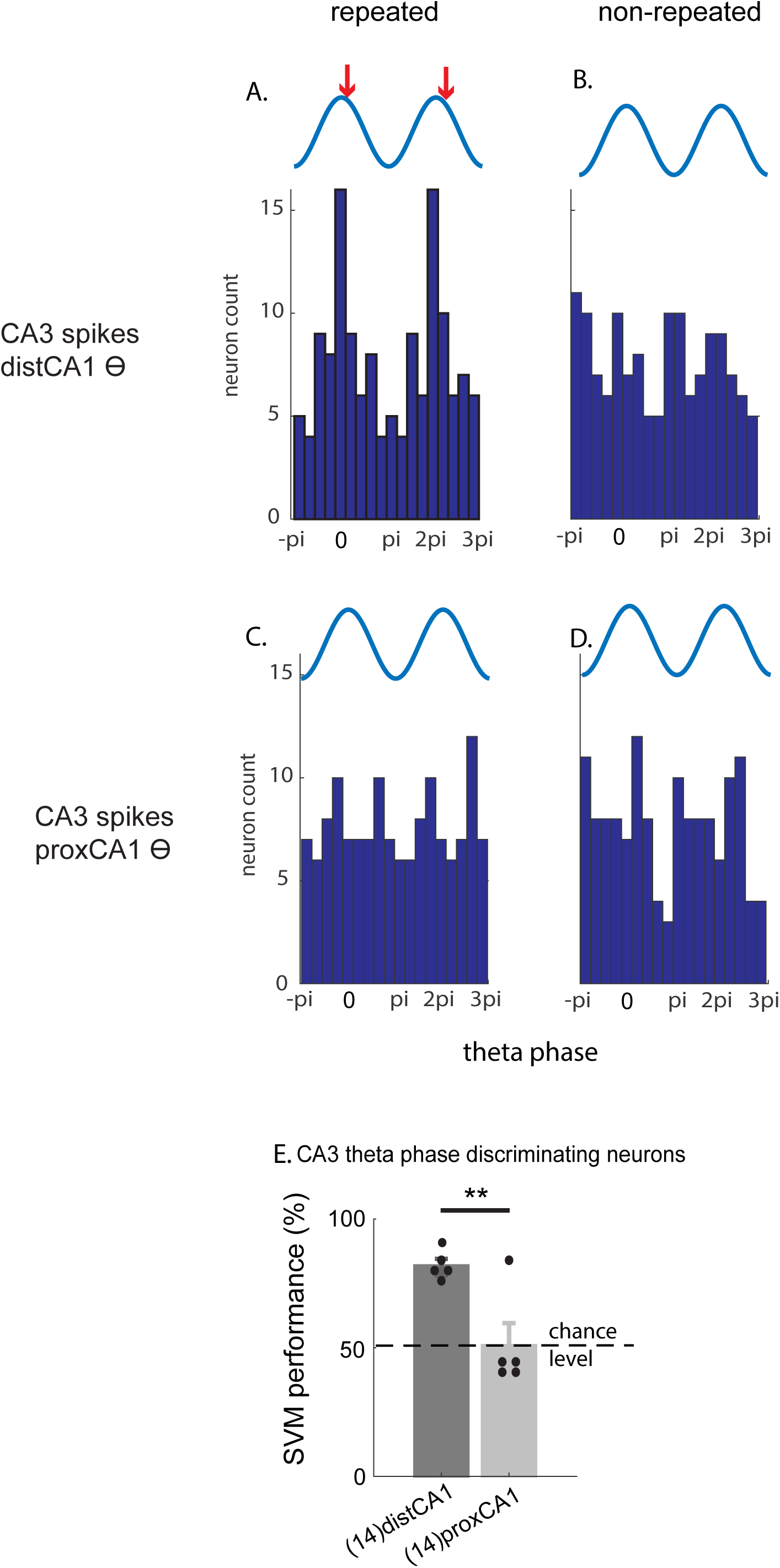
CA3 phase-discriminating neurons are preferentially phase-locked close to the peak of distal CA1 theta for the retrieval of experienced (‘repeated’) odors and classification performance of population CA3 spikes-distal CA1 theta phases predicts memory performance. The population distribution of the preferred angle of the theta phases of discriminating neurons across the whole CA3 region is concentrated close to the peak of theta for the retrieval of the memory of ‘repeated’ odors when distal CA1 theta is used as a reference (A, mean angle = 0.2198, var =0.7575) but not during the presentation of ‘non-repeated’ odors (B, var = 0.9516). No preferred angle is identified when theta in proximal CA1 is used as a reference for either ‘repeated’ (C, var=0.9734) or ‘non-repeated’ (D, var =0.8984) odors. E) SVM classification performances using theta phases of CA3 spikes either referencing to distal CA1 or proximal CA1. The classification performance using the phases of the CA3 neurons referencing to distal CA1, but not to proximal CA1 theta, is above chance level and comparable to memory performance. This indicates that interregional coupling (spike to theta oscillations) between CA3 and CA1 can predict memory performance trial-by-trial and plays an important role in recognition memory, especially between CA3 and distal CA1 for odor memory. The number of CA3 theta phase discriminating neurons per animal is in bracket. CA3 neurons are recorded along the entire proximodistal axis of CA3. dots: individual SVM performance; error bars: S.E.M. ** p< 0.01. Red arrows: mean angle of the population cell firing.

This result was confirmed by a SVM population analysis showing that the phase-information of CA3 discriminating neurons referenced to distal CA1 theta could predict better memory performance trial-by-trial than the phase-information referenced to proximal CA1 (SVM performance: distCA1: 89%, proxCA1: 49% correct; *t*-tests to chance level: *p* =.00023 and *p* =0.5, respectively; 2-sample t-test between distal and proximal CA1 theta : t(8)=3.4659=, p=0.0085; Fig. 6E). Hence, these results establish that the phase-locking of population CA3 spikes to distal CA1 theta plays a preponderant role in supporting odor memory retrieval trial-by-trial when compared to theta-phase locking to proximal CA1.

#### d) Local spike-theta phase-locking in CA3 or CA1 is less critical for memory retrieval

Subsequently, we evaluated to which extent the coordination of the spiking activity with local theta oscillations in CA3 or CA1 at the population level might contribute to memory retrieval.

In CA3, most neurons spiking during the retrieval of both ‘non-repeated’ and ‘repeated’ odor memory were phase-locked to theta in proximal CA3 (90%, 135/150, 27 neurons per animal) or distal CA3 (90%, 121/134, 24.4 neurons per animal). 51% of the neurons phase-locked to proximal CA3 (69/135, 13.8 neurons per animal) were phase discriminating neurons while 47% neurons (57/121, 11.4 neurons per animal) were phase-locked to distal CA3. No preferred angles were found for the phase-locking of CA3 spikes to proximal CA3 or distal CA3 theta oscillations for either odor type (proxCA3: ‘repeated’: p=0.257, ‘non-repeated’: p=0.2654, Fig 7A,B; distCA3: ‘repeated’: p = 0.8809, ‘non-repeated’: p = .272; circular Rayleigh tests, Supp.Fig. 10A,B) and SVM classification performances for CA3 discriminating neurons phase-locked to proximal or distal CA3 theta were low and did not differ significantly from chance level (proxCA3 theta: 67% correct, *p* = .0673, Fig. 7E; distCA3 theta: 46.25% correct, p=0.391, Supp. Fig. 10E; *t*-tests to chance level, Bonferroni corrected). These results suggest overall that local spikes-theta phase-locking in CA3 does not contribute in a critical manner to memory performance in this task.

**Figure 7.**
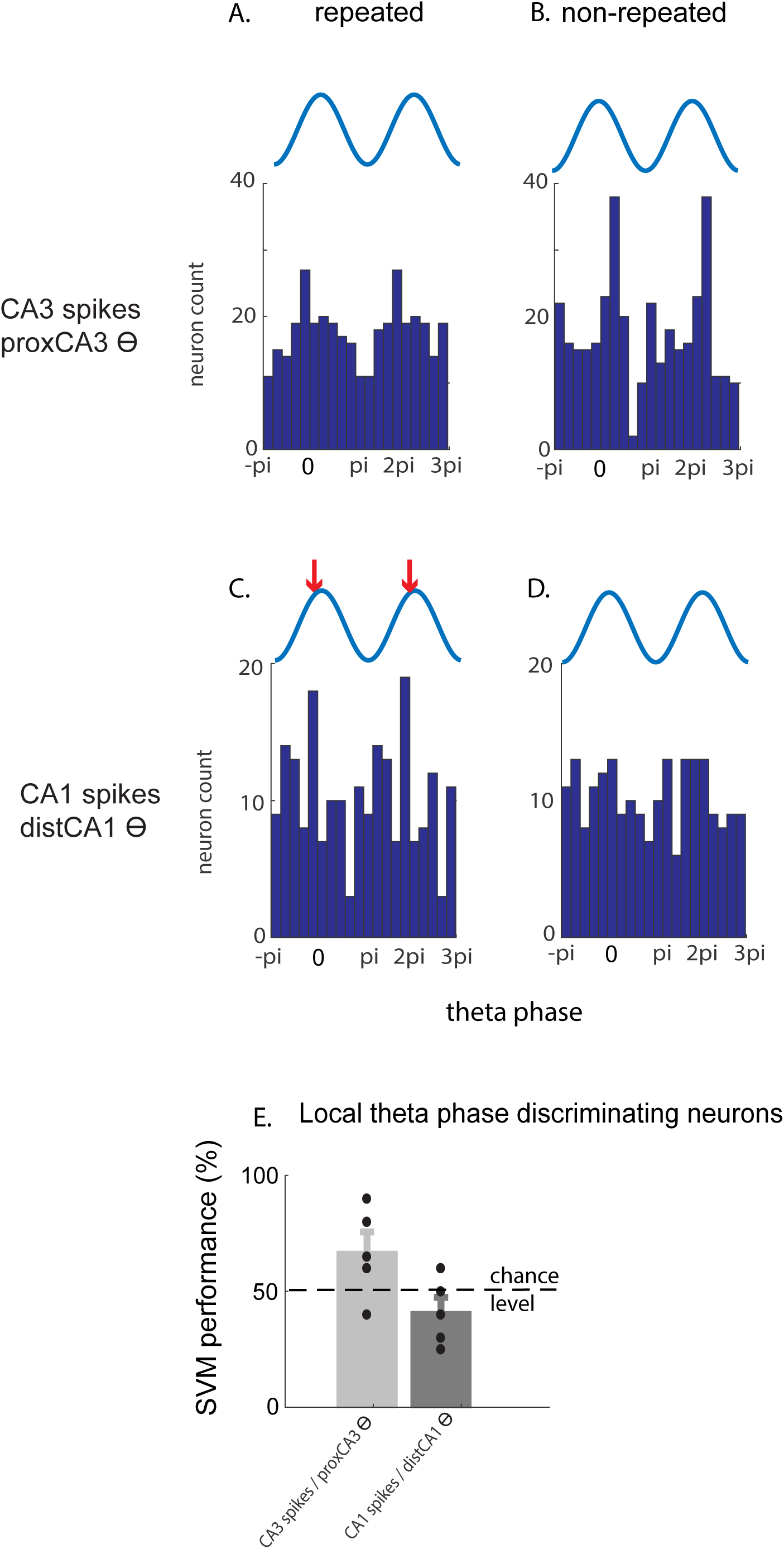
Local spike-theta phase locking in CA1 or CA3 cannot account for the discrimination of ‘repeated’ versus ‘non-repeated’ odors trial by trial. A,B) Population distribution of the preferred angle of the phases of CA3 spikes for the presentation of ‘repeated’ (A, var=0.8595) and ‘non-repeated’ (B, var=0.8612) odors using proximal CA3 (proxCA3) theta as reference. No preferred angle could be identified for either stimulus type with proxCA3 as a reference. The population distribution of the phases of CA1 spikes is concentrated close to the peak when distal CA1 theta is used as a reference for the presentation of ‘repeated’ (C, mean angle =-0.1612, var=0.5453) but not ‘non-repeated’ odors (D, var=0.845). E) SVM classification performances using either the local CA3 or CA1 spike-theta phases. Neither local CA3 nor CA1 theta phases can predict memory performance above chance level (SVM performance do not differ from chance level). This suggests that intraregional coupling (spike to theta phase) in CA1 or CA3 does not contribute to odor recognition memory to a major extent. var: variance; dots: individual data; error bars: S.E.M; chance level: 50%. Red arrows: mean angle of the population cell firing.

Furthermore, in contrast to CA3, much fewer CA1 neurons were phase-locked to either distal or proximal CA1 theta oscillations: only 17% of the neurons spiking during both ‘repeated’ and ‘non-repeated’ odors retrieval were phase-locked to distal CA1 (41/239 neurons, 8.2 neurons per animal; against 17.6%, 42/239, 8.4 neurons per animal phase-locked to proximal CA1) with half of these neurons (21/41, 4.2 neurons per animal) being discriminating neurons (against 35.7%, 15/42, 3 neurons per animal for proximal CA1). Population CA1 spikes were selectively phase-locked close to the peak of distCA1 theta oscillations for the retrieval of repeated odors (‘repeated’: p=0.014, ‘non-repeated’: p=0.6093, circular Rayleigh tests, Bonferroni corrected, Fig. 7C-D) while no theta phase concentration could be observed when proxCA1 theta oscillations were used as a reference (Supp. Fig. 10C,D). However, the former phase-locking appeared not to be decisive for discriminating ‘repeated’ from ‘non-repeated’ odors, as the SVM performance of both populations did not differ from chance level (dist CA1 theta: 41% correct, p = 0.2326, Fig. 7E; prox CA1 theta: 44% correct: p = 0.3046, Supp. Fig. 10E; t-tests to chance level, Bonferroni corrected). Similarly to spike-theta phase-locking in CA3, these results indicate that local spike-theta phase-locking in CA1 did not contribute to a major extent to non-spatial memory performance in the present task. All together, these last results demonstrate that even when the spikes of a high proportion of discriminating neurons are phase-locked to their local theta (the case for CA3), the local coordination of these signals does not play an essential role in the discrimination between ‘repeated’ and ‘non-repeated’ odors trial-by-trial, further supporting the selective role of the interregional phase-locking of CA3 spikes to distal CA1 theta in non-spatial memory retrieval established in the previous paragraph.

## Discussion

Our results on theta spike timing in the hippocampus bring further insight into the cellular and network mechanisms underlying recognition memory in the hippocampus. We examined theta phase-locking of cells within and between CA3 and CA1 regions along the proximodistal axis of the hippocampus and its relationship to memory performance. To do so, we used support vector machine analyses and a rat version of a task with a high memory demand used in humans. We found that, theta power is high in the present odor memory task, as is the case for spatially motivated tasks, especially during the retrieval phase. In addition, we observed that the population phase-locking of CA3 spikes to CA1 theta, but not local coupling, predicts animals’memory performance which is indicative of an important role of the coordination of CA3 neuronal activity with CA1 theta oscillations in certain non-spatial memory tasks. Importantly, this phase-locking was observable only when theta was refered to the distal, but not the proximal part of the CA1 region, suggesting a functional proximodistal gradient along the CA1 axis. In addition, the findings of a higher proportion of ‘effective’ neurons in distal CA1 and proximal CA3 and the fact that only the combined population cell firing of these two areas accounts for memory performance, provide furher evidence that distal CA1 and proximal CA3 might play a more essential role for recognition memory when stimuli are non-spatial.

### CA3 spikes-CA1 theta coupling predicts successful memory retrieval

In the present study, we focused on interregional spike-theta coupling, specifically the spike to theta phase-locking between CA3 and CA1 and showed that the theta phase information of CA3 spikes predict animals’memory performance trial by trial when referenced to the distal but not the proximal part of CA1 in an odor recognition memory task with high memory load (Fig 6). These results complement a handful of studies that have shown intra-regional differences in spike-theta phase locking in CA1 between the retrieval of studied stimuli (i.e. ‘repeated’) and the retrieval of non-experienced (‘non-repeated’) stimuli using either spatial or non-spatial recognition memory tasks with low memory load such as foraging in environments or spontaneous object/odor recognition (Manns et al., 2007, Lever et al., 2010, Douchamps et al., 2013). We extended knowledge on this topic by investigating additionally interregional spike to theta phase locking between CA3 and CA1. Furthermore, the finding that CA3 spikes preferentially phase-lock to CA1 theta at the peak of the cycles during memory retrieval is in line with the prediction of a prevalent computational model according to which CA3 inputs should be the strongest at the peak of CA1 theta during retrieval (Hasselmo et al., 2002, Manns et al., 2007). Direct empirical evidence for a preferential phase-locking of CA3 neurons to the peak of CA1 theta was however missing. Here, we show that CA3-CA1 coupling takes indeed place shortly after the peak of the theta cycles, bringing clear *in-vivo* evidence that CA3 spiking activities and CA1 theta oscillations are coherent at the peak of theta during retrieval.

One striking finding in the present study is that the theta phase information of CA3 spikes predicts animals’ memory perfomance only when refered to the distal part of CA1 but not its proximal part. This proximodistal difference unlikely stems from ‘constitutive’ differences in theta oscillations’ amplitudes between proximal and distal parts of CA1 as no consistent pattern of a proximodistal gradient was found in the 5 naïve control rats recorded (i.e. untrained; Fig.4E). In contrast, theta power was enhanced in distal CA1 compared to proximal CA1 in all phases of the task as well as during freerun in rats trained on the task (Fig 4A-D, Supp. Table 7). A possible explanation for this enhancement of theta power in trained animals is a progressive tuning of distal CA1 to the processing of non-spatial (odors) information over the 2 months of training. Such a tuning, in terms of increased connectivity between functionally connected cells or brain areas, has been reported in previous studies in humans and rodents upon extensive training (Olofsson et al., 2019; Huang et al., 2016; Fuentes-Garcia et al., 2019; Li et al., 2014).

Further evidence for proximodistal gradients along the axis of CA1 and CA3 was found in trained rats in terms of the proportion of ‘effective’ neurons (i.e. neurons primarily contributing to memory performance in the task; Fig 2E,F and Methods for details) and in terms of population cell firing of neurons discriminating experienced odors (‘repeated’) from non-experienced ones (‘non-repeated’) (Fig.2B). According to these gradients, spiking activity of distal CA1 and proximal CA3 together predicts best memory performance. This finding of a preferential involvement of proximal CA3 and distal CA1 in odor memory retrieval, a type of non-spatial memory, is in line with recent electrophysiological and immediate-early gene studies reporting a primary contribution of proximal CA3 and distal CA1 to non-spatial (object and odor) information processing or a reduced spatial tuning, when compared to distal CA3 and proximal CA1 (Nakamura et al., 2013, Beer et al., 2018, Flasbeck et al., 2018, Henriksen et al., 2010, Burke et al., 2011, Vandrey et al., 2021). Importantly, distal CA1 receives heavy neuroanatomical projections from the lateral entorhinal cortex (LEC) and less inputs from the medial entorhinal cortex (MEC) in comparison to proximal CA1(Tamamaki and Nojyo, 1995, Naber et al., 2001, Fyhn et al., 2004, Hargreaves et al., 2005, Henriksen et al., 2010, Burke et al., 2011, Ito and Schuman, 2012, Li et al., 1994, Ishizuka et al., 1995, Amaral and Witter, 1989, Witter et al., 1989, Ishizuka et al., 1990) and these latter brain areas preferentially linked to non-spatial or spatial information processing, respectively (Save and Sargolini, 2017, Igarashi et al., 2014, Fyhn et al., 2004, Deshmukh and Knierim, 2011, Hargreaves et al., 2005). Moreover, unlike distal CA3 that receives rather homogenous projections from both blades of the dentate gyrus (DG), proximal CA3 is principally targeted by the exposed blade of the DG (exp. DG) that is not engaged by spatial experience (Claiborne et al., 1986, Chawla et al., 2005). Altogether, these results point towards the existence of a ‘non-spatial’ medial temporal lobe (MTL) subnetwork including the exposed blade of DG, proximal CA3, distal CA1 and the LEC that would preferentially deal with non-spatial information as opposed to spatial information (Nakamura et al., 2013, Beer et al., 2018). *In-vivo* evidence for such a network and for a functional interaction between areas belonging to this ‘non-spatial’ subnetwork was however missing. As a first step for the *in-vivo* study of this subnetwork, we report evidence of a preferential recruitment of proximal CA3 and distal CA1 in non-spatial memory retrieval. In addition, we demonstrate that these areas function as a network as CA3 spike-distal CA1 theta coupling predicts best memory performance in our task.

Importantly, it is well-accepted that CA1 theta is highly influenced and coherent with EC theta (Zutshi et al., 2022, Kamondi et al., 1998, Ormond and McNaughton, 2015, Schlesiger et al., 2015, Lopez-Madrona and Canals, 2021) and that CA3 receives stronger anatomical projections from the EC than from CA1 (Witter, 2007). Hence, it cannot be completely ruled that the phase-locking of CA3 spikes to CA1 theta might to some extent also reflects their coherence to EC theta. However, a gradient in theta power could be identified along the proximodistal axis of CA1 in the present study, and only the theta phase information of CA3 spikes using distal but not proximal CA1 theta could predict memory performance. This suggests that theta oscillations in CA1 might be modulated by the local circuitry to a larger extent than originally described (Grossberg, 2021, Lopez-Madrona and Canals, 2021), making it less probable that EC theta contributes to a major extent to the CA3 phase-locking findings reported here. In addition, the identification of a strong inhibitory projection from CA1 to CA3 (Jackson et al., 2014) and the report of a causal influence from CA1 to CA3 in a study with simultaneous spiking recording (Sandler et al., 2015) further support a possible entrainement of CA3 spikes by CA1 theta oscillations. In addition, other intra and extra-hippocampal theta oscillations propagated by volume conduction have been suggested to contribute to hippocampal theta oscillations (Makarov et al., 2010, Lęski et al., 2010, Schomburg et al., 2014, Lasztóczi and Klausberger, 2014) and the ‘rythmicity’ of these ocillations to differ across task phases during learning for some of these generators (López-Madrona et al., 2020). Hence, it also cannot be completely ruled out that volume conduction could contribute to the differences reported in the present study. However, our results show that the theta phase information of CA3 spikes using proximal CA1 or CA3 theta oscillations as a reference cannot predict memory performance (Fig 6E & 7E), suggesting that the propagation by volume conduction of theta oscillations emerging from these generators might not contribute to a major extent to these effects.

In contrast to the interregional spike-theta phase locking between CA3 and CA1, local coupling in CA1 or CA3 failed to reveal a strong relationship with memory performance, even though the local spike-theta phase-locking in CA1 or CA3 was found to be important in tasks involving spatial navigation (Dragoi and Buzsaki, 2006, Robbe and Buzsaki, 2009, Ito et al., 2018, Ito et al., 2015, Bourboulou et al., 2019, Yu and Frank, 2021, Fernandez-Ruiz et al., 2017, Oliva et al., 2016, Manns et al., 2007). Our results are however consistent with the handfull of studies using non-spatial stimuli reporting that even though most cells exhibit significant spike-theta phase locking locally, no significant relationship is found between local spike-theta phase locking and memory performance (Rangel et al., 2016, Sanders et al., 2019). Hence, our results suggest that intra and interregional spike-theta coherence might serve vastely different purposes, at least for odor recognition memory. Disentangling the role of intra and interregional spike-theta coherence in the hippocampus is beyond the scope of the present study and will require further independent investigations.

Finally, in line with recent studies showing that discrimination between stimuli was better predicted by population level analysis as opposed to individual cell firing (El-Gaby et al., 2021, Guzowski et al., 2004, Leutgeb et al., 2004), no specific patterns of increase or decrease firing rates was detected at the single cell level (Supp. Fig5). This together with our finding that the combination of population firing activities can discriminate repeated from non-repeated odors suggest that differences in population firing activities between stimulus types might serve as an underlying mechanism for recognition memory rather than a general enhancement or suppression of cell firing rates.

Altogether, these results bring compelling *in-vivo* evidence that the coordination between CA3 neuronal activity and CA1 theta oscillations as well as population cell firing play important roles in recognition memory and predict memory performance in a path specific manner (i.e. by engaging preferentially the distal part of CA1 and proximal CA3 for non-spatial recognition memory). Moreover, these findings complement previous theoretical and empirical studies by providing clear evidence that the population CA3 neurons do fire close to the peak of CA1 theta cycles during successful recognition memory. In addition, our results show that neither local spike-theta coupling nor individual cell firing made a significant contribution within this frame.

### Theta oscillations are enhanced during non-spatial memory

Plethora of studies investigating spatial navigation and spatial memory have reported modulations of theta oscillations in CA1 (Whishaw and Vanderwolf, 1973, Wyble et al., 2004, Terrazas et al., 2005, Kunz et al., 2019, Hoffmann et al., 2015, Petersen and Buzsaki, 2020), while much fewer have examined the role of theta range oscillations in memory tasks devoid of salient spatial components (Pacheco Estefan et al., 2021, Leszczynski et al., 2015, Axmacher et al., 2010, Canolty et al., 2006). Therefore, we put an emphasis in the present study on the fact that high theta power is elicited during a non-spatial memory task and modulated by the different epochs of the task. Indeed, our results show that theta power is high during all phases of the task, but is especially strong at retrieval (Fig3 A-B).

Our result of an enhanced theta power in CA1 during a non-spatial memory task is in line with the findings of one of the very few studies investigating non-spatial memory in rodents, which however focused on intraregional couplings (i.e. did not investigate interregional couplings as is the case in our study) and used different behavioral tasks (spontaneous object recognition memory and odor-cued delayed nonmatch-to sample tasks; Manns et al., 2007). Our data bring also further support to studies in humans reporting a similar pattern in the hippocampus ‘as a whole’(Cohen et al., 2015, Klimesch, 1999, Vulic et al., 2021, Kragel et al., 2021, Tambini et al., 2018, Hebscher et al., 2021, Pacheco Estefan et al., 2021), showing for example that theta oscillations reflect cognitive demands in non-spatial tasks by possibly organizing neural processes through traveling theta waves (Zhang et al., 2018).

The increase in theta power observed in CA1 during memory retrieval in our task unlikely stems from an increased in locomotor activity as reported during running in previous studies (Korotkova et al., 2018, McFarland et al., 1975, Buzsaki et al., 1981, Buzsaki et al., 1985, Buzsaki and Moser, 2013, McClain et al., 2019) as rats’ behavior during recording bouts is comparable across study and test phases of the task: rats rest during the baseline and are virtually stationary for the 2 sec bouts following stimulus delivery during the study or the test phase (speed <5 cm/sec during any phase of the task). Of note, such a low speed is in the range of the speeds typically excluded from studies reporting a dependency of theta amplitudes and frequency on running speed (range of speeds typically considered: 5 to 100 cm/sec; see for examples: Kaefer et al., 2020, Stella et al., 2019). Hence, this might partially explain why our results do not support a strong dependency of theta oscillations’ amplitudes on running speed.

Besides running: stress, arousal and alertness levels might also enhance theta oscillations (Mikulovic et al., 2018, Hata et al., 1987, Yamamoto, 1998, Chang, 1992, Nollet et al., 2019, Murthy et al., 2019, Shors et al., 1997, Liberman et al., 2009). Stress and arousal levels are expected to be fairly low in our task given that animals are heavily handled and trained daily for over 2 months. In addition, alertness levels are likely to be comparable during the study and the test phases of the task as, in both cases, rats are left undisturbed 20 min prior to study and test (which corresponds to the baseline and delay period, respectively). Hence, differences in stress, arousal or alertness levels between epochs are unlikely to account for the differences in theta power observed between retrieval and other phases of the task in CA1 (baseline and study). Thus, our results together with the full breath of the literature on spatial memory and navigation, point toward a robust role of CA1 theta oscillations in memory function that is independent of the spatial content of the memory.

Overall, our results show that theta oscillations play an important role in non-spatial memory, as is the case for spatial navigation and memory. In addition, we demonstrate that the coordination of population firing in CA3 with CA1 theta oscillations (spike-theta phase locking) at retrieval occurs close to the peak of theta only for stimuli for which a memory could be formed (‘repeated’ stimuli) and can account for memory performance. This indicates that interregional synchronization of neuronal spiking with theta oscillations might facilitate discriminative processes tied to memory function, especially between CA3 and distal CA1 in the case of non-spatial memory. In contrast, similar coupling at the local level in CA1 or CA3 as well as rate or phase coding at the single cell level appear to play less of a decisive role in recognition memory.

## Author Contributions

S.K. and M.S. designed the experiments; S.K. and E.A. conducted the experiments; S.K. and N.A. analyzed the data with J.C. and M.S.’s feedback; S.K. and M.S. wrote and J.C. and M.Y edited the manuscript. M.S. is corresponding for the conceptual and mechanistic framework underlying the design of the experiments; S.K. is corresponding for formulating and identifying the electrophysiological mechanisms.

## Acknowledgements

We would like to thank J. Maiwald for her assistance in animal behaviour training, experiments and brain slice preparation; D. Koch for her assistance in recording drive building and brain slicing; H. Mulla-Osman for her assistance in setting up DeepLabCut Analysis; Dr. K. Kaefer and J. Wallenschus (IST Austria) for their initial technical support snf Dr. S. Mikulovich for her comments on an early version of the manuscript; Dr. C. Reichert for his comments on SVM analyses. This project is funded by the DFG CRC 779 and 1436.

---

**Supplementary Figure 1.**
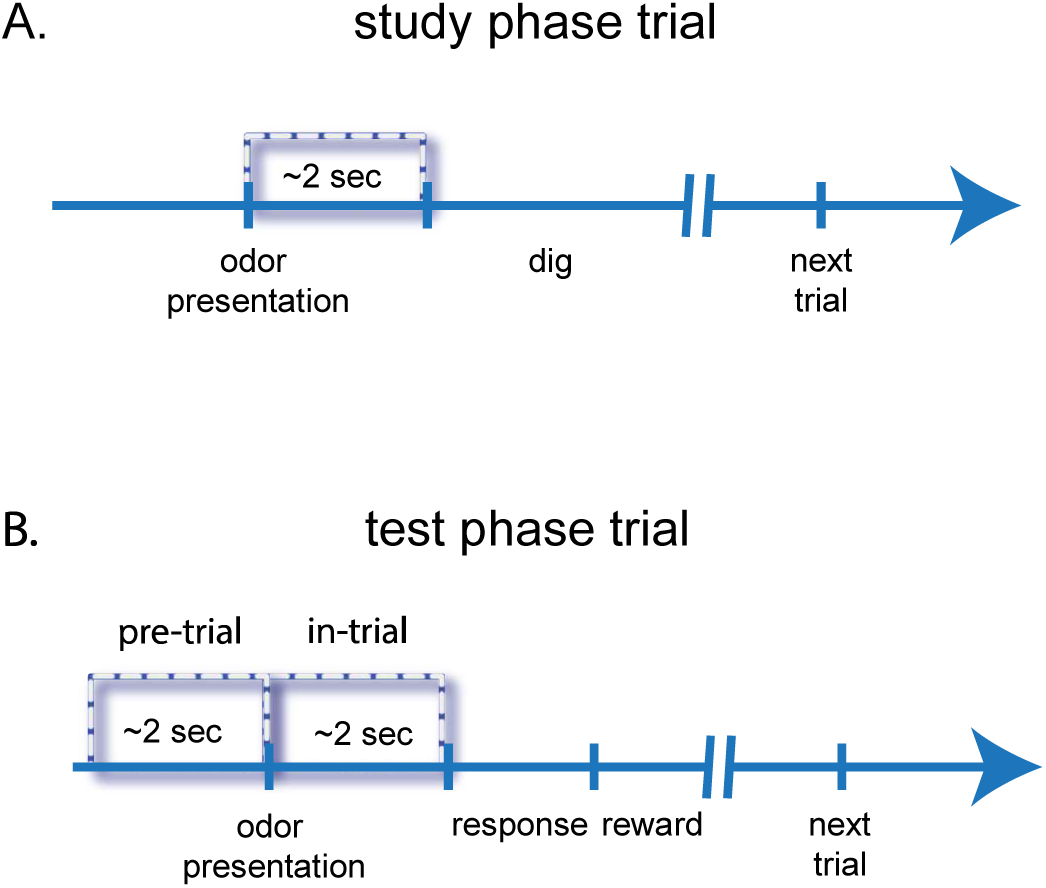
Time-windows for analyses for a ‘study’ trial and a ‘test’ trial: electrophysiological analyses were performed during the 2 sec following stimulus delivery for the study phase and 2sec before or after stimulus delivery (‘pre’ and ‘in-trial’ periods) for the test phase. The time-window of 2 sec was defined based on prior experiments showing that behavioral response in this task occurred shortly after 2 sec.

**Supplementary Figure 2.**
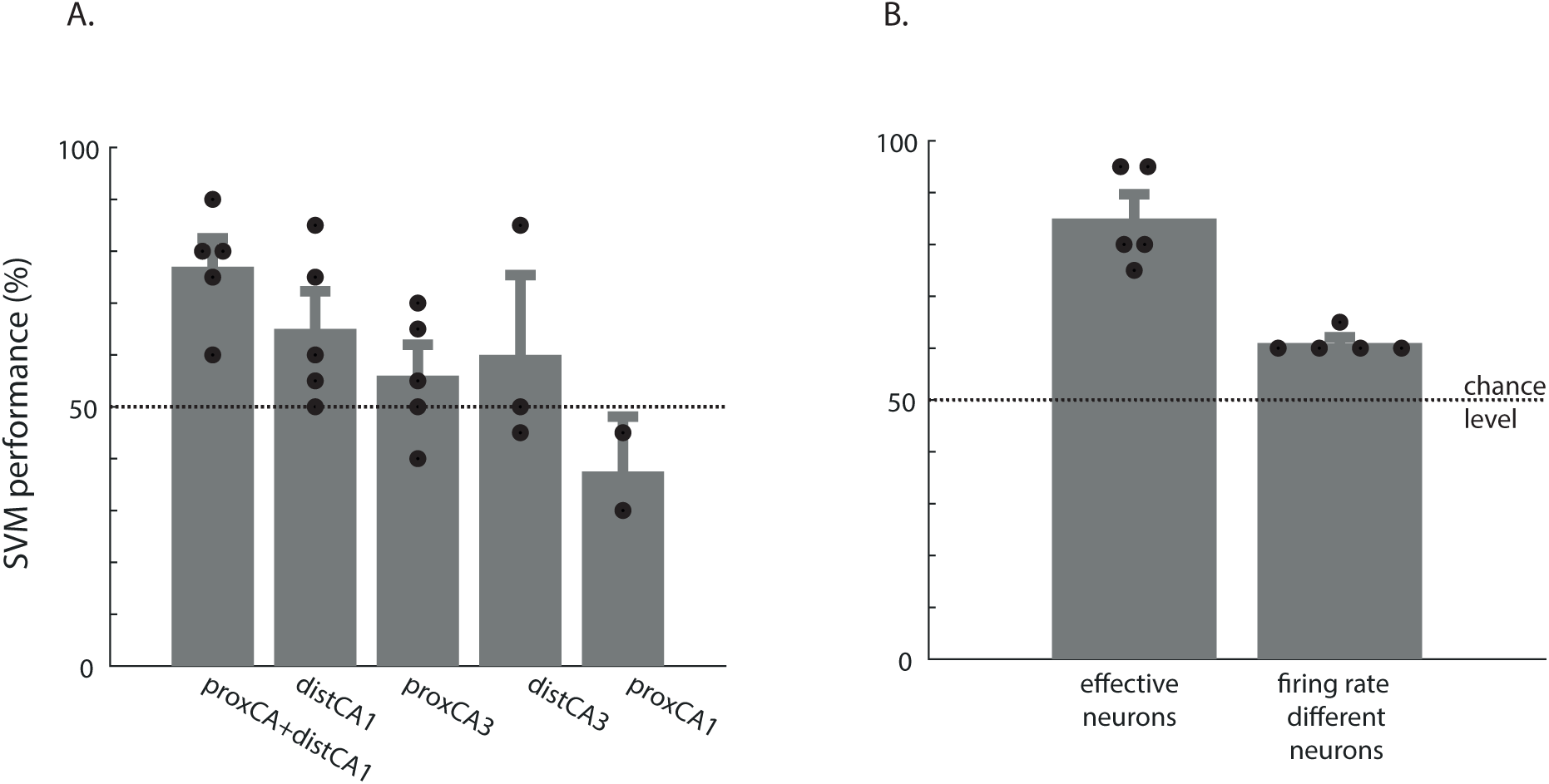
SVM performances using the population firing activities of neuronal populations recorded in different hippocampal subregions (A) as well as that of effective neurons or neurons which differ in firing rate for repeated and non-repeated odors (B). Only SVM performances proxCA3+distCA1 and effective neurons differ from chance level (both ps <0.0053). dots: individual SVM performance; error bars: S.E.M ; chance level: 50%.

**Supplementary Figure 3.**
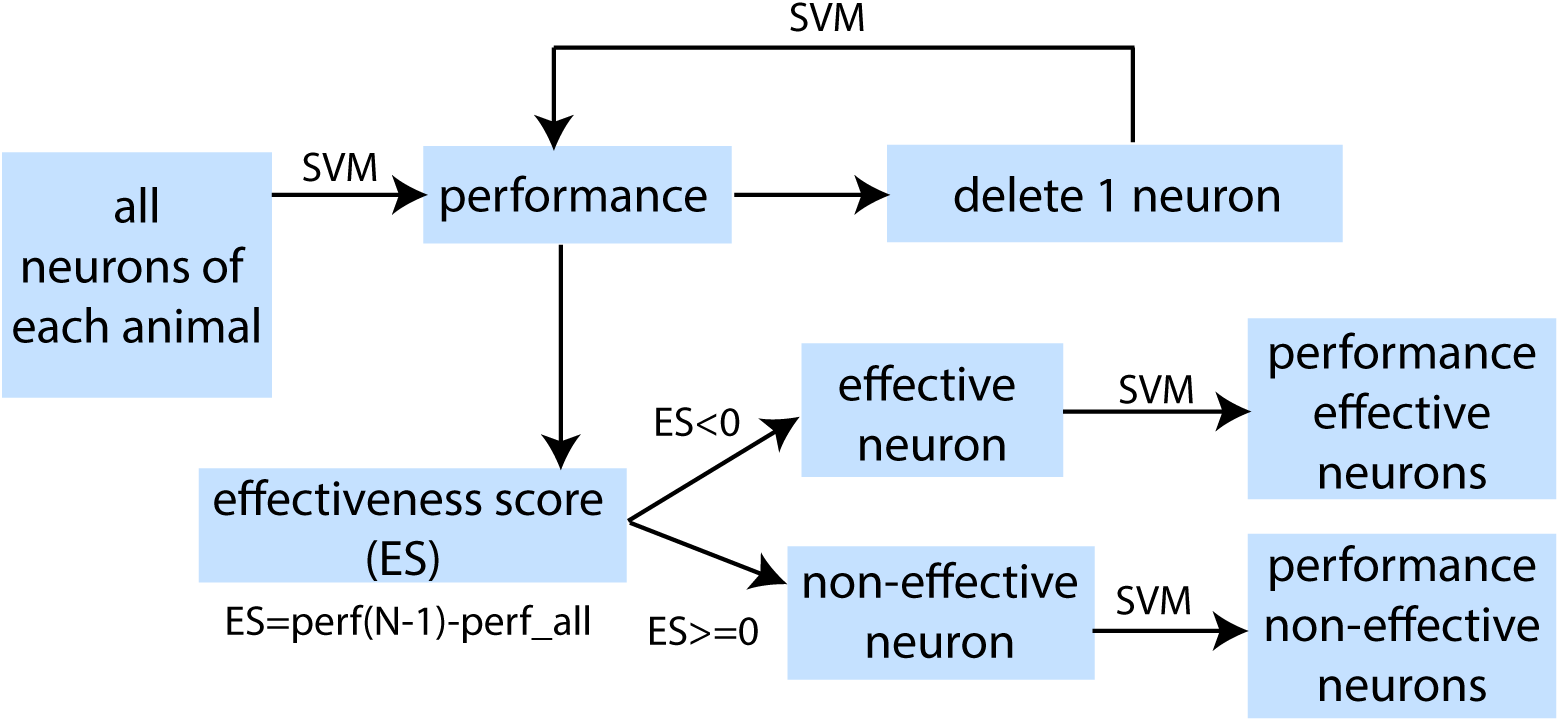
Schema of the iterative feature selection procedure for categorizing individual neurons into ‘effective’ or ‘non-effective’ neurons using Support Vector Machine. (Ku et al., 2008, Brinkmann et al., 2015). In brief, for each iteration, each neuron was deleted one at a time from the pool of recorded neurons (n=N). The effectiveness score (ES) was calculated by subtracting the performance of all recorded neurons (perf all, n=N) from the performance of the remaining neurons (perf(N-1), n=N-1) to assess if the deletion of one specific neuron affected the population classification performance. If that was the case (ES<thr; threshold), the neuron was termed ‘effective neuron’, if not: ‘non-effective’ neuron. Once tested, each deleted neuron was placed back to the pool of neurons, and the next iteration started with the deletion of the next neuron (without repeating sampling).

**Supplementary Figure 4.**
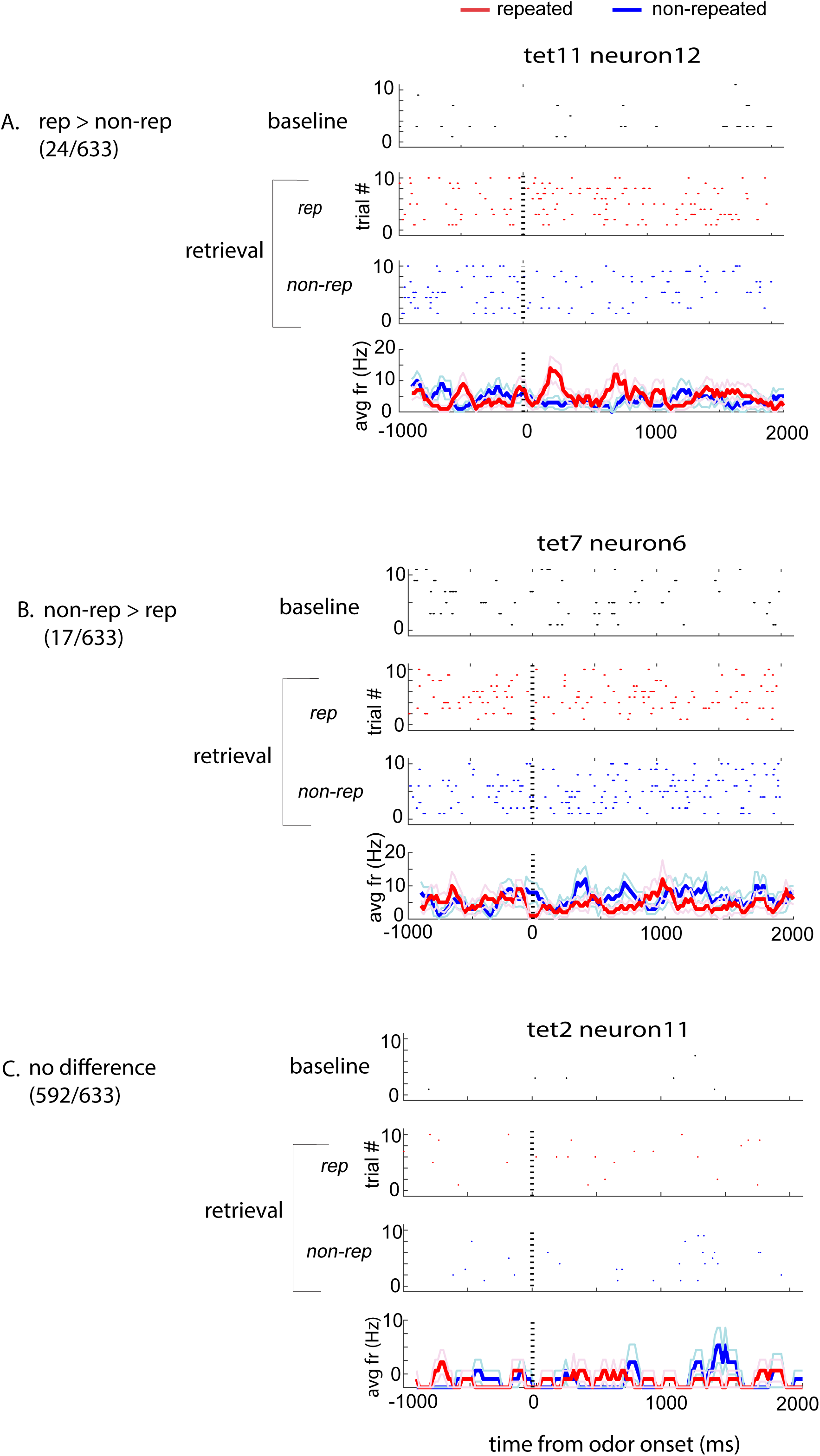
Single cell firing patterns of representative neurons during baseline and retrieval. Examples of firing rates: higher for ‘repeated’ than ‘non-repeated’ odors A), higher for ‘non-repeated’ than ‘repeated’ odors B) or not differing between stimulus types C). The three first plots of each panel are trial-by-trial spike raster plots. The forth plot shows averaged firing rates across 10 trials with S.E.M. (SEM indicated by lighter colored lines). Dashed vertical lines indicate the onset of the odor presentation.

**Supplementary Figure 5.**
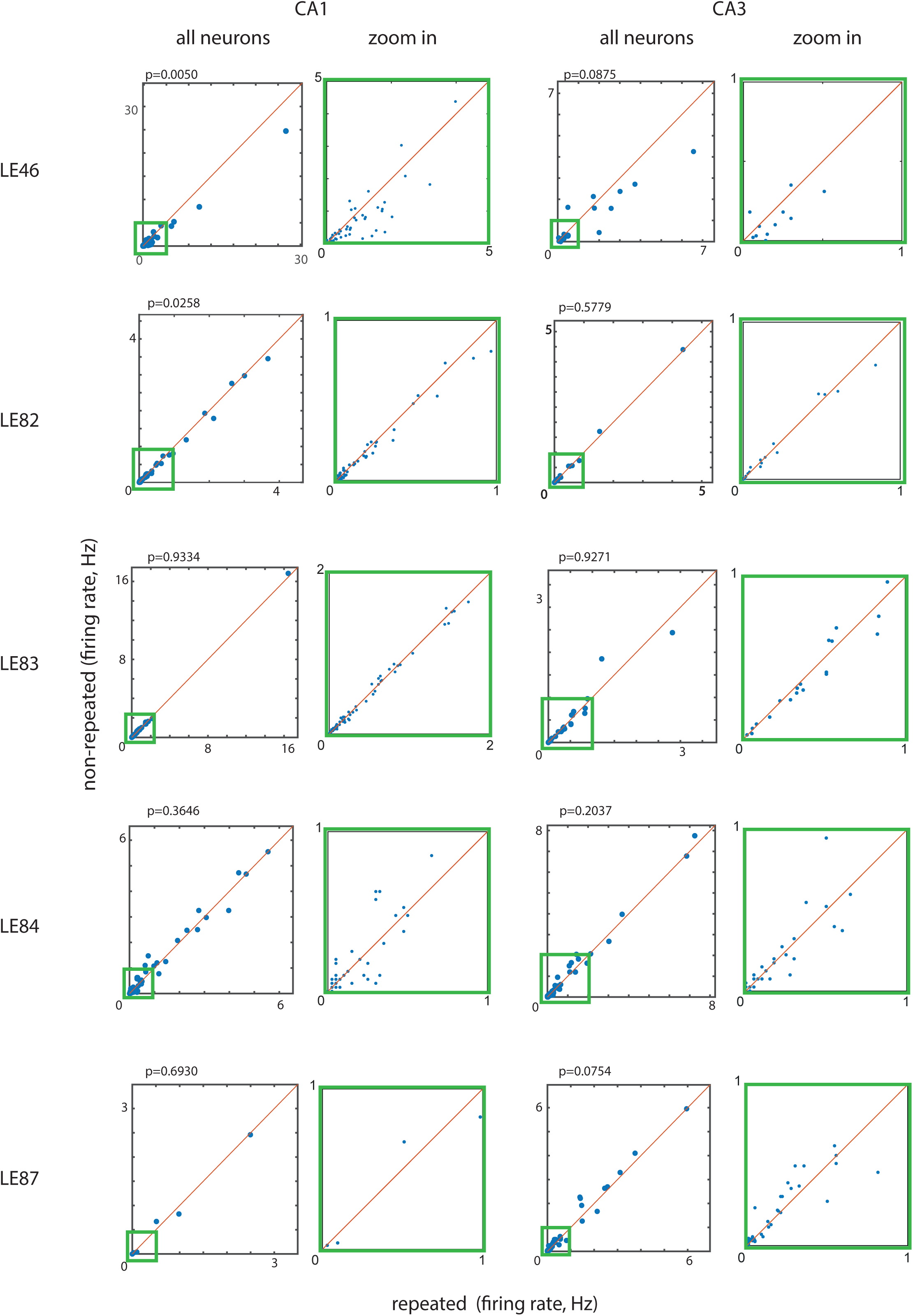
Mean firing rates of all recorded neurons do not show clear differential patterns for “repeated” or “non-repeated” odors. The firing rate patterns were not consistently different at test in either CA1 or CA3 upon ‘repeated’ or ‘non-repeated’ odor presentation. Mean firing rate was higher when rats smelled ‘repeated’ than ‘non-repeated’ odors only in two out of five animals (LE46, LE82) in CA1 while no significant differences could be observed in the three remaining rats (LE83, LE84, LE87). In CA3, firing rates do not differ beween ‘repeated’ or ‘non-repeated’ odors for any of the animals recorded. Blue dots: individual neurons. For CA1 and CA3, a zoom-in of the left side plot is shown on the right side.

**Supplementary Figure 6.**
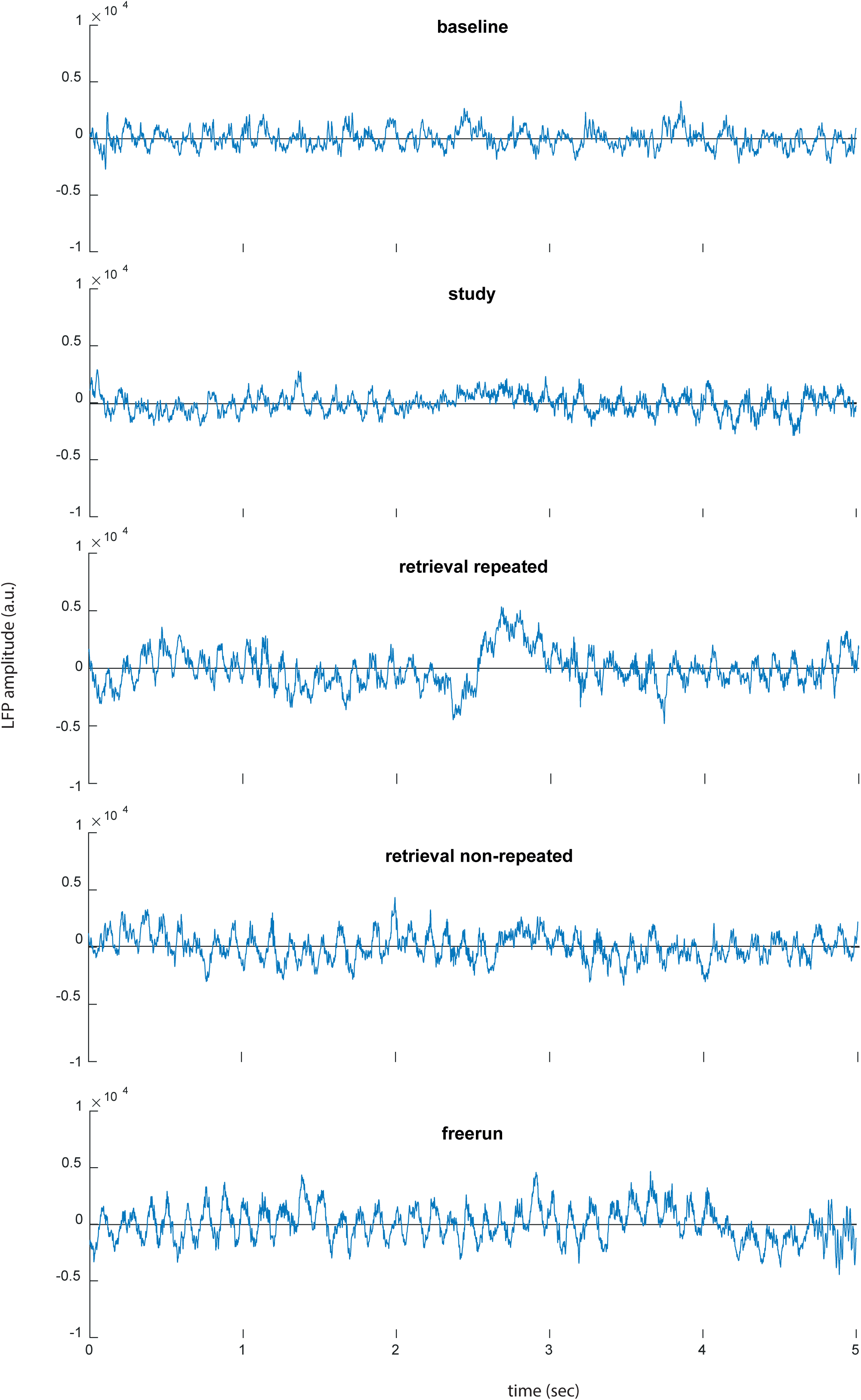
Example of 5-second long raw local field potential (LFP) traces of one representative animal.

**Supplementary Figure 7.**
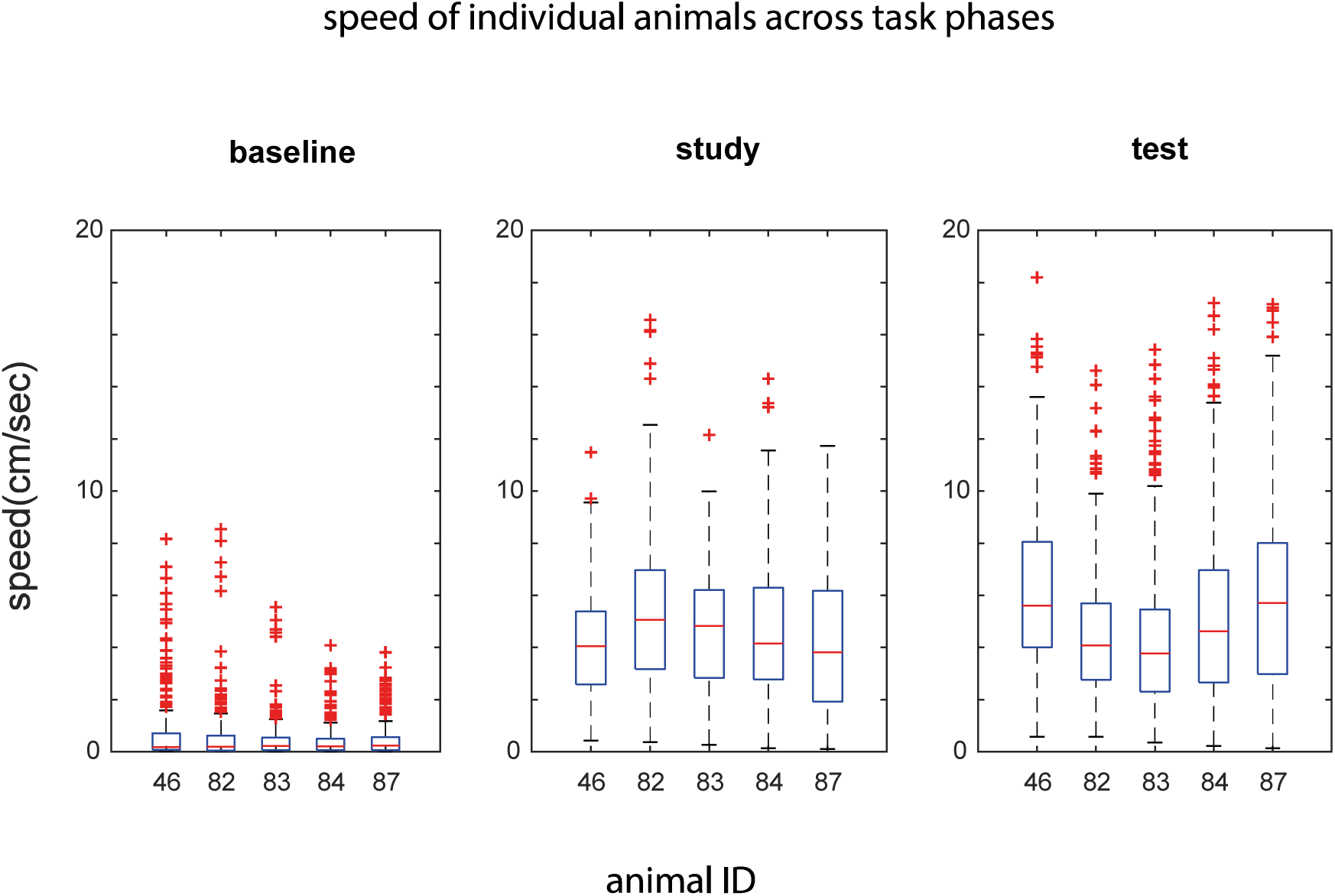
Speed of individual animals across different task phases. Speed at baseline is very low (average speed: 1.35cm/sec) and significanly lower than during study and test (4.34 and 4.69, respectively, ps < 0.001, 2-sample t-test). Of note, speed at study and test are also quite low and comparable to speed excluded in tasks reporting a dependency of theta amplitudes on animals’ speed (i.e. <5cm/sec). Central marks (red): speed medians, the bottom and top edges (blue): 25 and 75 percentiles; whiskers: one times the interquantile range, +: outliers.

**Supplementary Figure 8.**
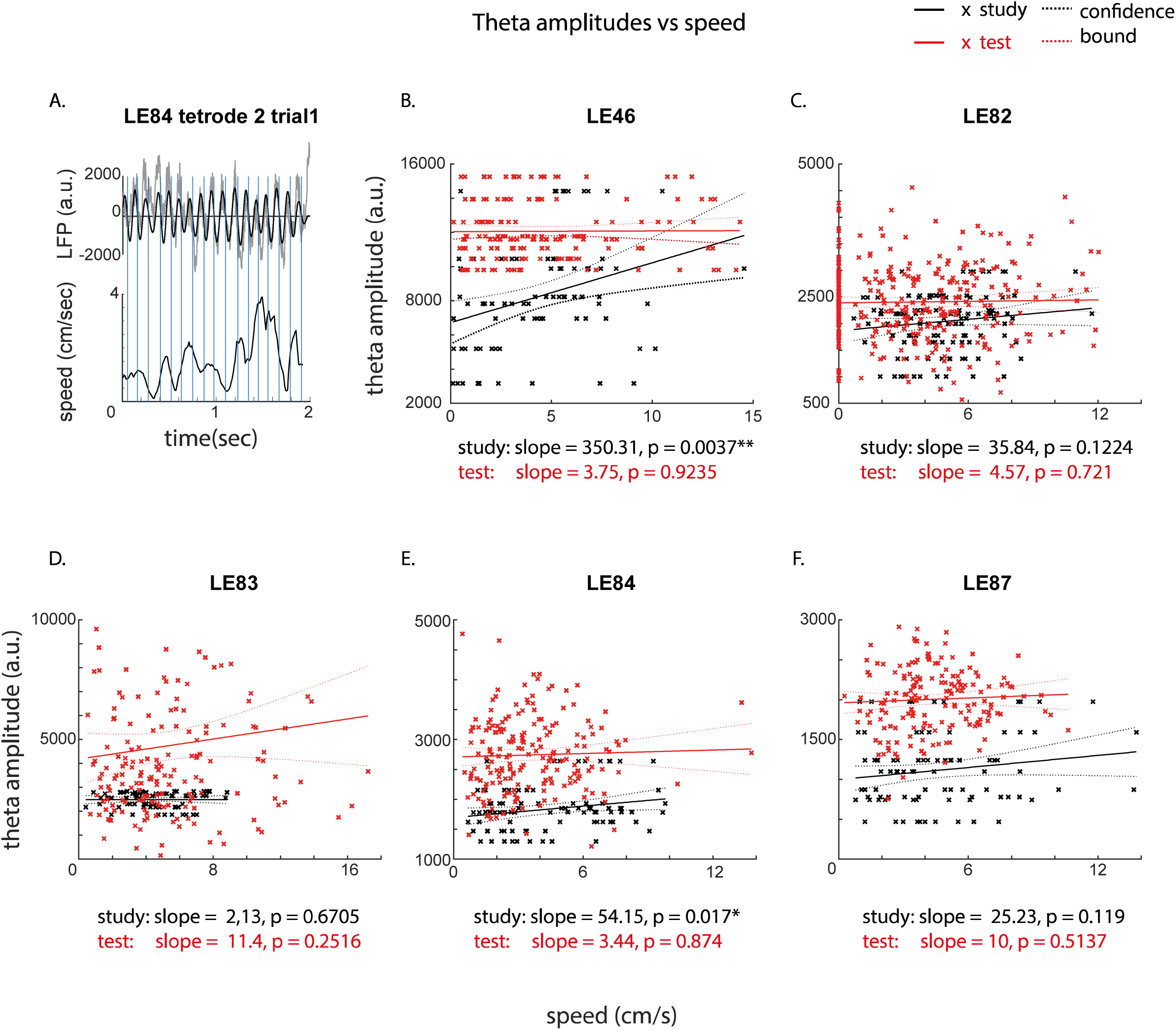
Analysis of theta amplitudes and speed during study and test for each animal, showing no clear dependency between theta amplitudes and speed in the present task. (A) The local field potential (LFP, grey) overlaid with filtered theta cycles (6-12Hz, black, top panel) and the speed of one representative animal in a test trial. Blue vertical lines indicate the middle points of the peak and the trough of each theta cycle and their corresponding speed. (B-F) For a direct comparison, the theta amplitude of each cycle was plotted as a function of speed estimated by averaging the speed during a 80ms time-window, centering at the middle of the peak and the trough of each theta cycle (blue verticle lines in A). A linear regression and a least square approach were applied to test for the dependency between theta amplitudes and speed. For each animal, the study (black) and the test (red) phase are fitted with an independent model. Theta amplitudes did not vary in function of the speed of the animals at test for all animals nor during study for 3 out of the 5 recorded rats (but see LE46 and LE84), indicating a general lack of robust dependency between theta amplitudes and speed in the present task. Note that the average speed of animals in this task is very low and comparable to the range of speeds excluded from studies reporting a strong dependency between theta power and animals’ speed (exclusion criterium: 5cm/sec). X: amplitude of each theta cycle and its corresponding speed; solid line: linear fit; dotted line: 95% confidence bound. *:p<0.05; **: p<0.01.

**Supplementary Figure 9.**
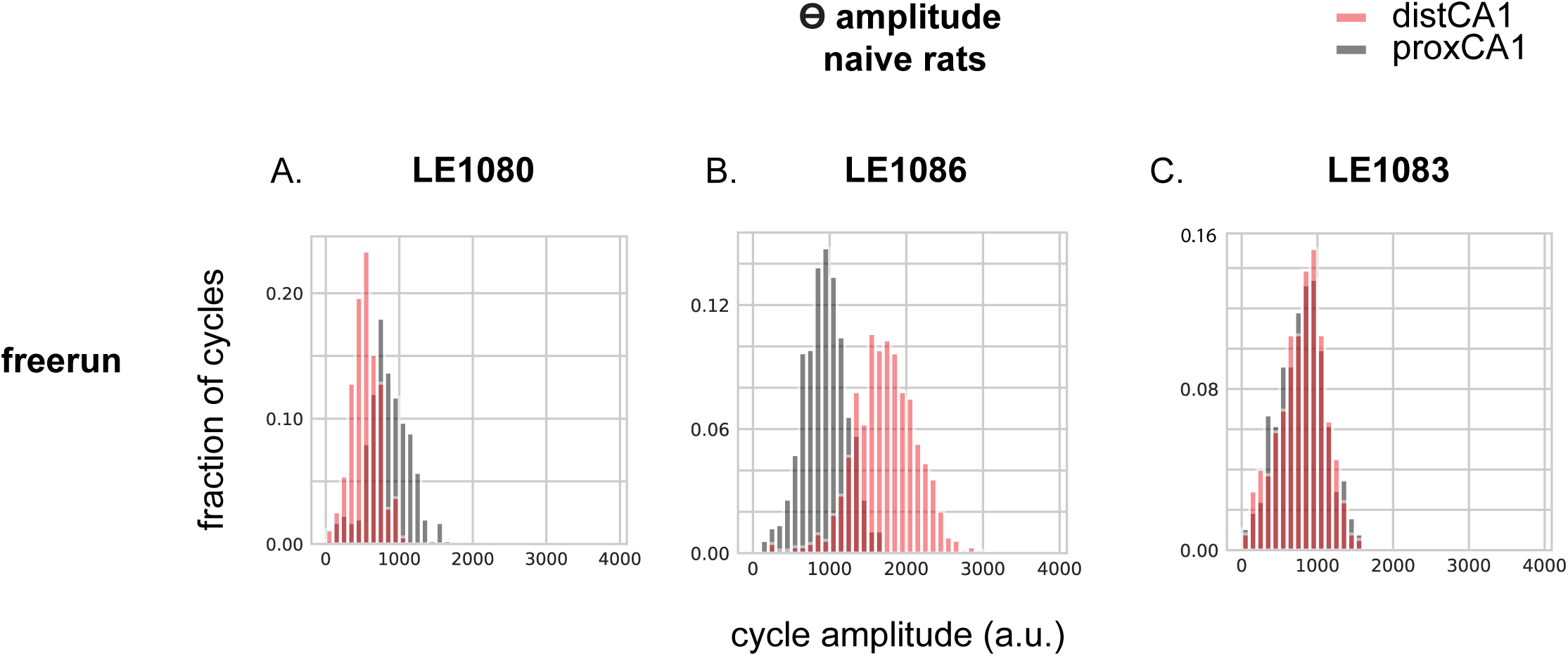
Representative examples of distributions of theta amplitude in distal CA1 and proximal CA1 in naïve rats during freerun. Out of the five naïve rats recorded, two showed higher theta cycle amplitudes in proximal CA1 (A); two others higher in distal CA1 (B) and no differences was observed between distal and proximal CA1 in the last rat (only one example shown per category). The inconsistent distribution among naïve rats indicate a lack of coherent ‘constitutive’ difference in theta power along the proximodistal axis of CA1.

**Supplementary Figure 10.**
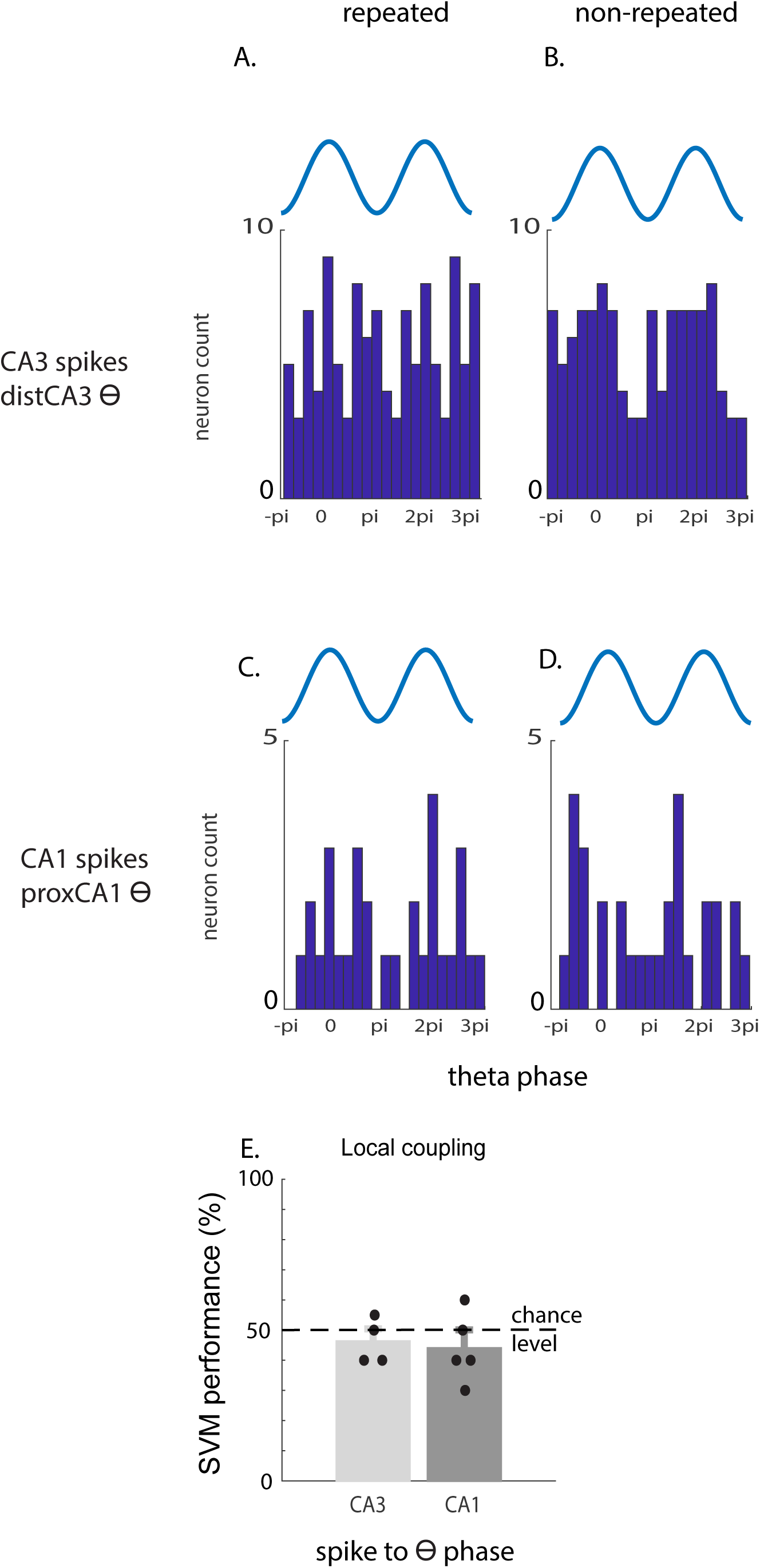
As seen in Figure 7 for local spikes-theta phase-locking in distal CA1 or proximal CA3, local couplings to either distal CA3 theta (A-B) or proximal CA1 theta (C-D) can not account for memory performance. A-B) Theta-phase locked CA3 neurons did not fire at a preferred angle of distal CA3 theta for either ‘repeated’ A) or ‘non-repeated’ odors B) at test. C,D) Similar results were obtained for the phase-locking of CA1 neurons using proximal CA1 theta as reference for the presentation of ‘repeated’ (C) or ‘non-repeated’ odors (D) respectively. E) Population classification performances using either one of these populations were not different from chance level, indicating that local theta phase-locking in CA1 or CA3 does not contribute significantly to recognition memory performance. Dots: individual SVM performance; Error bars: S.E.M.

## Methods

### Surgery

Once animals learned the task (see ‘Behavioral paradigm and training’ section) an electrode microdrive housing 16 individually movable tetrodes was implanted (Axona Ltd, St. Albans, UK, http://www.axona.com/). Tetrodes were arranged in 3 rows targeting the right dorsal hippocampus and covering the whole proximodistal axis of CA1 and CA3 from AP -3,3 to AP -4mm. Tetrodes, made of four tungsten wires (12 um diameter each, California Fine Wire Company, Grover Beach, CA), were twisted and heated to form a bundle. The tips of tetrodes were cut and gold-plated (Sifcoasc, OH, USA, Gold NC SPS 5355) to reach an impedance < 150kΩ with a NanoZ (Multichannel system, Reutlingen, BW, Germany). Animals were anesthetized by induction with 3% isoflurane and injected with Ketamin/Xylavet (75mg/kg and 5mg/kg) mixture and Atropine (0.1mg/kg) to facilitate breathing. Once deeply anesthetized, animals were placed in the stereotactic frame (David Kopf Instrument, CA, USA) fitted with a gas mask supplying 1% isoflurane and 0.8∼1.2L/min oxygen. Local anesthesia (Lidocaine, 0.5%, < 7mg/kg) was injected subcutaneously before incision. Analgesic agent Metacam (5mg/kg) was injected two hours after the injection of the Ketamine cocktail. A craniotomy was performed above the targeted hippocampal area (AP - 2.5∼-4.5, ML 1.2∼4.2, right hemisphere). Six anchoring stainless screws (1mm diameter, 3-4mm long) were placed into the skull above the prefrontal, left hemisphere and cerebellar areas. A silver wire was twisted around the two cerebellar screws to serve as ground electrodes and soldered onto the contact point of the Microdrive at a later time. After removing the dura using a dura hook (Fine Science Tools, Heidelberg, Germany) tetrodes were lowered to 800∼1000um below cortex surface. A mixture of melted paraffin wax/oil (wax:oil 3.5:1, Aldrich) was dropped along the tetrodes (covering the craniotomy), and the drive was secured with dental cement (Paladur, Heraeus, Hanau, Germany). Animals were supplied with oral analgesic meloxicam for 3 days and recovered for one week. Tetrodes were then slowly lowered to the regions of interest into the pyramidal layers of CA1 and CA3 over 2 weeks.

### Subjects and Stimuli

Adult male Long–Evans rats [350–450 g; *n* = 10 in total; were maintained under reverse light/dark cycle (7:00 A.M. light off/7:00 P.M. light on) at a minimum of 90% of normal body weight, handled a week before the experiment, and tested in their home cage. The stimulus odors were common household scents (thyme, paprika, coriander, etc.) mixed with playground sand. The scented sand was held in glass cups (one odor per cup), and a cup was fixed on a small platform lowered in the front part of the cage for testing. A pool of 40 household odors was available, 20 were used each day chosen in a pseudo random manner. All procedures were approved by the animal care committee of the State of Saxony-Anhalt (42502-2-1555 LIN) and performed in compliance with the guidelines of the European Community and ARRIVE guidelines.

### Behavioral paradigm and training

To test the non-spatial recognition memory, we used the innate ability of rats to dig and to discriminate between odors. Behavioral training and paradigm have been described in details previously (Ku et al., 2017; Mahnke et al., 2021; Nakamura et al., 2013). In short, each training session contained a “study” phase, a delay, and a “test” phase. Each day, rats were presented with a study list of 10 odors, which was different each day. Odors were chosen in a pseudo random manner. During the recognition, i.e. the “test” phase, animals were tested for their ability to distinguish between the 10 odors presented during the study phase (“repeated” odors) and 10 additional odors (“non-repeated” odors) that were part of the pool of forty odors, but were not presented during the study phase. To do so, rats were first trained to dig in the stimulus cups with unscented sand to retrieve one 1/4 of piece of loop cereal (Kellogg’s) after which animals were trained on a delayed non-matching to sample rule. During the “test” phase, when rats were presented with an odor that was part of the study list (a “repeated” odor), rats were required to refrain digging, turn around, and go to the back of the cage to receive a food reward: a correct response for a “repeated” odor (Fig. 1B; an incorrect response would be digging in the stimulus cup). Conversely, when the odor was not part of the study list (a “non-repeated” odor), animals could retrieve a buried reward by digging in the test cup: a correct response for a “non-repeated” odor (an incorrect response would be going to the back of the cage to receive the reward). To ensure that the task could not be solved by smelling the reward buried in the sand, crushed cereals were mixed in the sand of all stimulus cups. In addition, no spatial information useful to solve the task was available to the rats, given that testing cups for ‘repeated’ and ‘non-repeated’ ‘odors were presented at the exact same location. Reward locations differed for the ‘non-repeated’ and the ‘repeated’ odors (front and back of the cage, respectively), but were only experienced by the animals once a decision had been made (e.g., when the trial was over), hence could not contribute to behavioral performance. Training lasted for about 2 months and consisted of several steps, during which the number of studied odors increased from 1 to 10, the delay increased from 1 to 20 min, and the number of odors during the test phase increased from 2 to 20 (10 ‘repeated’ and 10 ‘non-repeated’ intermixed). Animals transitioned between successive training stages when performance reached a minimum of 75% correct for 3 consecutive days. Once this criterion was reached for the final training stage (10 study odors, 20 min delay, and 20 test odors), animals were further trained for at least another week and implanted with a microdrive. Following one week of recovery period, neurons were screened and animals trained back to criteria following which they were recorded for one additional session that was used for the analyses.

Animal’s local motion was minimized by using a 35×19 cm cage. The behavior of the animals was recorded and streamed to a computer with a high-speed, HD infrared camera (100fps, Genie Nano Series, GigE Vision, Teledyne Dalsa, Ontario, Canada). The camera sent one trigger per frame to the recording system to synchronize video and electrophysiological recordings. Behavioral performance was assessed with high temporal resolution (10 ms precision). Periods analyzed started upon stimulus presentation (when the front edge of the platform on which the stimulus cup was fixed crossed the wall of the cage, Supp. Fig.1) and ended two seconds later: i.e. for the ‘test’ phase: before the turning or digging behavioral responses. The time-window of 2 sec was defined based on prior experiments showing that behavioral responses in this task occurred shortly after 2 sec. Rats were sniffing and alertly waiting during these periods.

For trained rats, free-run was performed in an 80×80×10cm square arena (open top, customized) following the delayed non-matching to odor task, after animals rested for 20 min in their home-cage. The arena was placed in the recording booth and animals could foraged for cereals (cheerios) scattered on the floor. A freerun session lasted 8 minutes. Behavioral and electrophysiological signals were recorded with the camera and recording systems used while animals performed the task. The same freerun procedure applied to naïve controls, except that animals did not undergo the two-month training on the task, but were housed in the same facility and brought to the experimental room as frequently as experimental animals. Naive animals were implanted with the same recording drive (16 tetrodes) as trained animals and recorded after one week of recovery as was the case for trained animals. The recording session for naïve rats lasted about 30 minutes and consisted of 20 minutes resting time (home cage) and 8 min freerun.

### Histology and Reconstruction of Recording Positions

After the final recording, animals were overdosed with pentobarbital (500mg/ml, IP) before intracardiac perfusion with 0.9% saline and 4% formaldehyde. The brain was extracted and post-fixed with 4% formaldehyde for 24hrs. Brains were then transferred to a 30% sucrose solution for cryoprotection and subsequently frozen at -41°C in isopentane using dry ice and stored in -80°C. Brains were cut into 30um coronal sections with a cryostat (Leica CM3050S, Leica Microsystems), mounted on glass slides and stained with cresyl violet (Nissl staining). The tips of the tetrodes were identified from the stained brain slices.

### Data Acquisition

The electrophysiological signals are preamplified with headstages (2×32 channels, either Axona Ltd, St. Albans, UK or Neuralynx, Bozeman, MT, USA) then digitized at either 24kHz (Axona Ltd, St. Albans, UK, animal LE46) or at 32kHz (Neuralynx, Bozeman, MT, USA, animals LE82, E83, LE84, LE87 and control animals) with a 64 channel data acquisition system. Broad band and low-pass filtered (<4800Hz) data are simultaneously recorded with the Axona system. Broadband and clustered data (default) are simultaneously recorded with Neuralynx.

### Quantification and Statistical Analysis

#### Behavior

##### Memory performance

The performance of animals (% correct choice) were calculated by evaluating the number of correct trials during the test period divided by all testing trials (20) and multiplying by 100.

##### Speed analysis

To ensure that theta amplitude differences between task periods do not solely stem from animals’ motion, the speed of the animals was assessed using DeepLabCut software package (Mathis et al., 2018). Because of the similarity in behavior across animals (n=5), ‘study’ and ‘test’ video clips were pooled together to train one network. For the same reason, baseline video clips of trained (n=5) and naïve (n=5) animals were also pooled together. Freerun videos were pooled to train a third independent network. 50 frames per video were randomly extracted, and the nose, ears and 3 points along the midline of the body (neck, mid of the body and tail base) were manually labeled on each frame. The networks were trained with default parameters and 200 000 iterations. The trained networks and frames were then manually evaluated to analyze the entire videos.

Given that the body of the animals was virtually immobile during the present task, the position of the mid-point of the two ears was used to estimate the speed of the animals in an attempt to prevent an under estimation of the speed of movements. The instantaneous moving speed was estimated based on the displacement of the labels occurring between two consecutive frames, to which a Kalman filter was applied (Fyhn et al., 2004). The speed of individual animals was calculated by pooling the 2-second trials together and the median of the speed distribution was used to represent the speed because of the non-normal distribution of the speed across the task phases. For baseline and freerun periods, 20 periods of 2 seconds each were randomly selected and the same procedure applied.

#### Spike Sorting

The raw data (sampled at 24kHz or 32kHz) were filtered between 500 and 8000 Hz and an automatic clustering procedure applied using klustakwik (http://klusta-team.github.io/klustakwik/). The parameters are as follows: action potentials with power of > 4 standard deviations from the baseline mean were selected and their spike features were extracted with principle component (PC) analysis. Using the first 3 PCs and the initial cluster was set to 100. Afterwards the clusters were manually selected, combined or split with the software klusters (http://klusta-team.github.io/klustakwik/) or M-clust (https://redishlab.umn.edu/mclust), selecting only the clusters with clear refractory periods in their autocorrelation, well-defined boundaries and stability over time.

#### Support Vector Machine (SVM)

The performances for classifying the two testing trial types (‘repeated’ or ‘non-repeated’ odors during test period) with either the population spiking activities or the theta phases of population CA3 or CA1 spikes were assessed with the support vector machine method (SVM method; Ambard and Rotter, 2012; Ku et al., 2008) using software LIBSVM https://www.csie.ntu.edu.tw/~cjlin/libsvm/index.html). A polynomial kernel was used with default parameters.

The leave-1-out cross validation procedure was used to evaluate the accuracy of SVM classification performances. In brief, for each iteration, the population neuronal activities of one trial was left out of the dataset and the data of the rest of the trials (19 out of 20) were used to train the SVM. The trained SVM was then tested with the trial left out in the current iteration. The same procedure was performed for each trial without repetition. Finally, The SVM performance was calculated by counting how many iterations yielded a correct output. Since the test phase of the task consists of 10 ‘repeated’ and 10 ‘non-repeated’ trials, chance level mount to 50% correct. Given the total number of test trials, a minimum reduction in performance is 5% (1/20 trials).

For population spiking activities, the combined mean firing rates of the 2 sec following stimulus presentation during retrieval phase of each neuron were used as ‘features’. To test whether the hippocampal population firing predicted the type of stimulus presented (‘repeated’ vs. ‘non-repeated’) or behavioral responses (‘digging’ or ‘turning’ independently of whether animals were correct or not), the SVM were trained separately with the labels ‘stimulus types’ or ‘behavioral responses’. For population theta phases, the circular mean of the theta phases of each 2-sec trial was calculated for each neuron and the circular means of the theta phases of all neurons per trial used to perform the SVM-training and the cross validation. Of note, for theta phase analyses, only the neurons which spiked for each of the 20 test trials were used. Neurons which did not fire at each trial were excluded since the mean theta phase could not be calculated for all trials.

#### Selection of ‘effective’ neurons

To find out which neuron contributed “effectively” to the SVM performance, i.e. which neurons would cause a significant decrease of SVM performance if removed, an iterative feature selection procedure was applied to the neurons of each animal (Supp. Fig. 3). For each iteration, each neuron was deleted one at a time from the pool of recorded neurons (n=N). The effectiveness score (ES) was calculated by subtracting the performance of all recorded neurons (perf all, n=N) from the performance of the remaining neurons (perf(N-1), n=N-1) to assess if the deletion of one specific neuron affected the population classification performance. If that was the case (ES< threshold), the neuron was termed ‘effective’ neuron, if not: ‘non-effective’ neuron. Once tested, each deleted neuron was placed back to the pool of neurons, and the next iteration started with the deletion of the next neuron (without repeating sampling). The procedure was illustrated in Supp. Fig. 3.

To evaluate the ES threshold, the features of the 20 trials were bootstrapped 1000 times and a 5%-decrease in SVM performance was found to be significant with 95% confidence interval, hence set as a (ES) threshold. To further ensure that removing one effective neuron caused a significant reduction in SVM performance, the dataset of the target/studied neuron was replaced by randomly selecting 20 data points from the remaining N-1 neurons (i.e. out of 20 X (N-1) data points), the SVM analysis was performed with this new dataset (in total N neurons) and this procedure repeated 1000 times. Doing so, we found that the SVM performance of this new dataset (n=N) was comparable to the performance of the N-1 dataset (i.e. 5% lower than the performance of all neurons) despite the same sample size (n=N), indicating that the drop in performance did not stem from the sample size alone and that removing the ‘effective’ neuron significantly affects the SVM performance.

To evaluate whether the distribution of ‘effective’ neurons is homogeneous along the proximodistal axis of CA1 or CA3 and compare across animals, the recording sites of ‘effective’ and ‘non-effective’ neurons was sorted into 5 CA1 and CA3 segments for each animal and the count of neurons were summed across animals for each segment (Fig. 2D, sup. Table 2). The proportion of ‘effective’ neurons was calculated by dividing the sum of ‘effective’ neurons by the total number of neurons recorded (effective + non-effective across animals) for each segment and a linear regression model with a least square approach was fitted to the CA1 and CA3 data.

#### Theta Oscillation Analyses

The local field potential (LFP) recorded with the Axona system (4800Hz) were directly used for further analysis; the LFP recorded with the Neuralynx system was low-pass filtered with cutoff frequency = 300Hz using Matlab filter function then down-sampled 8 times to 4k Hz. Toolbox Chronux (version 2.12 v03) was used to obtain the theta power and to estimate the theta phases using multi-taper method (taper=5-9). The cycle-by-cycle theta amplitudes were obtained with the toolbox bycycle (https://github.com/bycycle-tools/bycycle, (Cole and Voytek, 2019)). Theta frequency was defined as oscillations between 6-12 Hz.

For each animal, theta cycle amplitudes of all trials (10 to 20 cycles/trial) in a specific task phase were pooled together (see Figure 3C-D and Figure 4 for distributions in a representative animal), subsequently to which a group analysis on trained (N=5) and naïve animals (N=5) was performed. Population differences in theta amplitudes were calculated using the Kolmogorov–Smirnov (KS) test to generate D values (D=Maximum|Fo(X)−Fr(X)|, X=theta cycle amplitudes, F(X)=observed cumulative frequency distribution of a random sample of n observations) and Friedman’s test was performed on these values to obtain group statistics.

To test whether the theta phases of neuronal spiking can discriminate different stimulus types (‘repeated’ vs ‘non-repeated’ odor), theta phases of each neuron were pooled together for ‘repeated’ or ‘non-repeated’ trials and compared with the circular Watson-Willians multi-sample test for equal variance (threshold p<0.05, Matlab Circular Statistic toolbox, Philipp Berens 2023, https://www.mathworks.com/matlabcentral/fileexchange/10676-circular-statistics-toolbox-directional-statistics). When the difference in theta phases between ‘repeated’ and ‘non-repeated’ trials surpassed the threshold (p=0.05, 2-sided), neurons were termed ‘phase-discriminating’; when it did not, neurons were termed ‘non-phase discriminating’. Non-phase discriminating neurons were subdivided into two categories: type I neurons, phase-locked to the same angle of CA1 theta oscillations for both ‘repeated’ and ‘non-repeated’ odors and type II neurons, phase-locked to only one stimulus type.

##### Statistical Analysis

Analyses were performed using custom written Matlab (R2020a) or python scripts (version 3.8). The normality of all distributions were tested with Lilliefors test. When data were normally distributed, ANOVA and one-or two-tailed t-tests were used. For non-normal distributions (theta amplitudes in distCA1 or proxCA1), the Mann-Whitney U-test was used for unpaired data, the Wilcoxon signed-rank test for paired data or comparisons to chance level and the Kruskal-Wallis test for more than 2 groups. The significance threshold of all tests were Bonferroni corrected: for SVM performances better than chance for different hippocampal subregions: threshold: p=0.05/5=0.01 (Fig. 2B); for comparing ‘effective’ to ‘non-effective’ neurons: threshold: p=0.05/3=0.017 (Fig. 2C); for post-hoc comparison of theta amplitudes between baseline, study and test phases: threshold: p=0.05/3=0.017; for comparing animals’ speed: threshold: p=0.05/2=0.025; for testing the theta-phase information using SVM: threshold: p=0.05/2=0.025 (Fig. 6E and Fig. 7E). Statistical tests used and p values are reported in the text, tables or figure legends. (*p<0.05, **p,0.01, ***p<0.0001). Error bars indicate the S.E.M.

##### Data and Code Availability

Codes used for the present study will be available upon publication on github. Data will be available upon request.

---

**Supplementary table 1.** Population classification performances and cell counts of each experimental animal

|  |  | #LE46 | #LE82 | #LE83 | #LE84 | #LE87 | average | stats |
| --- | --- | --- | --- | --- | --- | --- | --- | --- |
| proxCA3 | cell no. | 19 | 11 | 22 | 18 | 29 | 19,8 | t(4)=1,12 |
|  | SVM perf (%) | 65 | 40 | 55 | 70 | 50 | 56 | p=0,32 |
| distCA3 | cell no. | 0 | 49 | 2 | 49 | 29 | 25,8 | t(3)=1,12 |
|  | SVM perf (%) |  | 45 |  | 85 | 50 | 60 | p=0,34 |
| proxCA1 | cell no. | 25 | 11 | 0 | 20 | 14 | 14 | t(1)=-1,67 |
|  | SVM perf (%) | 45 |  |  | 30 |  | 37,5 | p=0,34 |
| distCA1 | cell no. | 47 | 110 | 76 | 71 | 20 | 64,8 | t(4)=2,3 |
|  | SVM perf (%) | 75 | 85 | 60 | 55 | 50 | 65 | p=0,08 |
| firing rate | cell no. | 11 | 8 | 6 | 18 | 2 | 9 | t(4)=11 |
| diff. neurons | SVM perf (%) | 65 | 60 | 60 | 60 | 60 | 61 | p=1,94E-14 |

Abbreviations:
proxCA3:proximal CA3; distCA3: distal CA3; proxCA1: proximal CA1; distCA1. distal CA1; perf: performance; no: number
Stats: one-tailed t-test comparing the SVM performance to chance level (50%)
Firing rate diff neurons: neurons with significantly different firing rates between repeated and non-repeated trial, see supp. Table 3
Effective neurons vs. Firing rate diff. neurons: ttest for pairwise comparison t(4)=5,2372, p=0.0064
The distribution of the pairwise difference between effective and firing rate different neurons is normal
See supp. Fig2 for bar plots.

**Supplementary Table2.** Distribution of effective neurons in each segment and animal

| cell<br>count | #LE46 |  | #LE82 |  | #LE83 |  | #LE84 |  | #LE87 |  | sum |  | proportion(%)<br>(effective/all) |
| --- | --- | --- | --- | --- | --- | --- | --- | --- | --- | --- | --- | --- | --- |
|  | effective | total | effective | total | effective | total | effective | total | effective | total | effective | total |  |
| **CA1 segment** |  |  |  |  |  |  |  |  |  |  |  |  |  |
| 1 |  | 0 | 5 | 22 | 2 | 15 | 5 | 26 |  | 0 | 7 | 63 | 11,11 |
| 2 |  | 11 |  | 27 | 1 | 17 | 3 | 25 | 3 | 6 | 7 | 86 | 8,14 |
| 3 | 4 | 36 | 4 | 61 | 3 | 30 |  | 20 |  | 14 | 11 | 161 | 6,83 |
| 4 | 1 | 13 | 1 | 4 | 2 | 14 | 1 | 20 |  | 14 | 5 | 65 | 7,69 |
| 5 | 1 | 12 |  | 7 |  | 0 |  | 0 |  | 0 | 1 | 19 | 5,26 |
| **CA3 segment** |  |  |  |  |  |  |  |  |  |  |  |  |  |
| 1 |  | 0 | 6 | 11 | 2 | 6 |  | 0 |  | 7 | 8 | 24 | 33,33 |
| 2 | 1 | 19 |  | 0 |  | 0 |  | 11 | 3 | 33 | 4 | 63 | 6,35 |
| 3 |  | 0 |  | 13 | 1 | 16 |  | 7 | 1 | 7 | 2 | 43 | 4,65 |
| 4 |  | 0 |  | 8 |  | 0 |  | 23 | 1 | 22 | 1 | 53 | 1,89 |
| 5 |  | 0 |  | 28 |  | 2 | 1 | 26 |  | 0 | 1 | 56 | 1,79 |
| sum | 7 | 91 | 16 | 181 | 11 | 100 | 10 | 158 | 8 | 103 | 47 | 633 | 7,42 |
| average per animal |  |  |  |  |  |  |  |  |  |  | 9,4 | 126,6 |  |

Effective: counting effective neurons in each segment of CA1 or CA3
Total: only neurons firing during test are considered.

**supplementary Table3.**
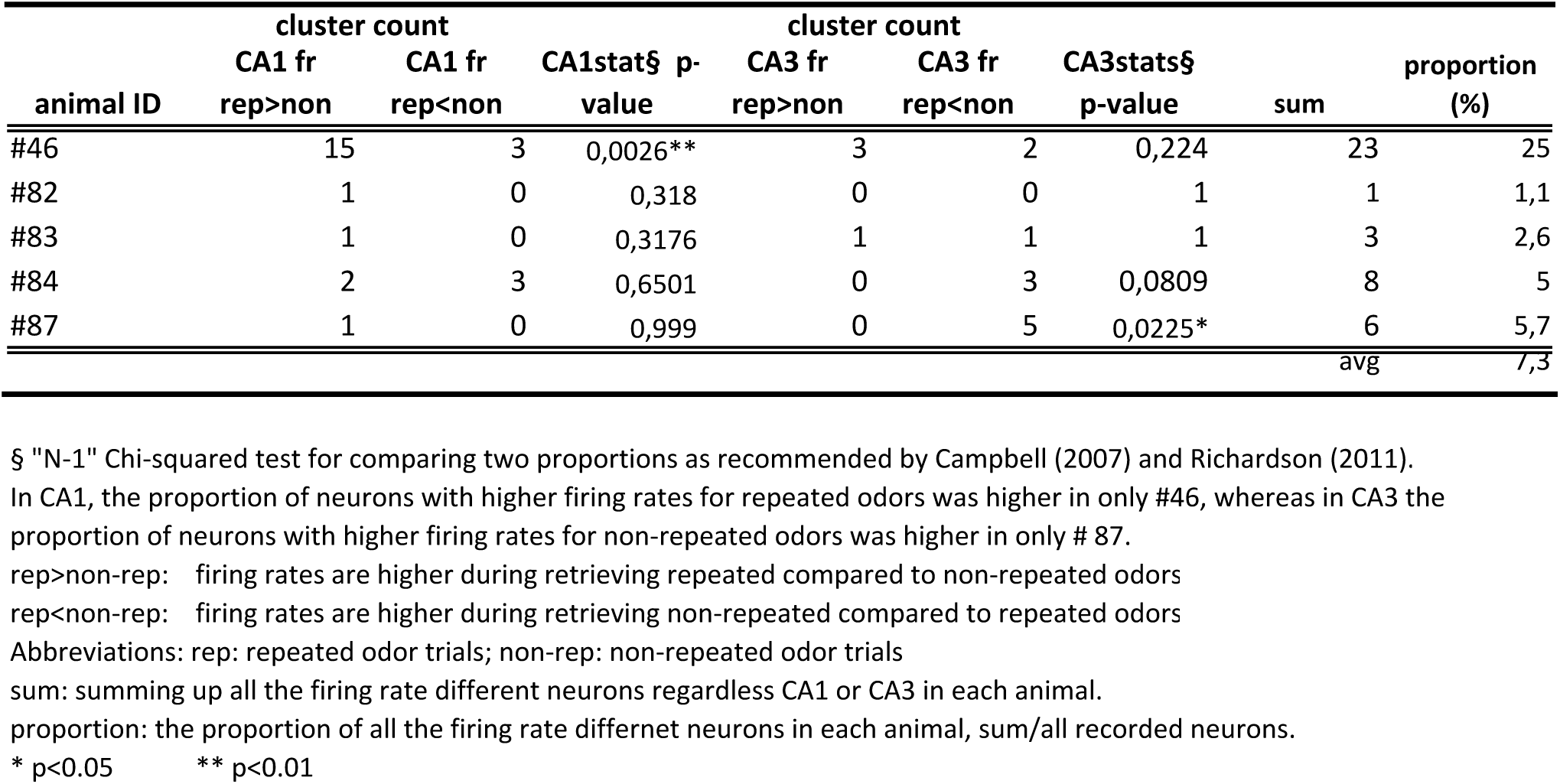
Firing rate difference analysis

**Supplementary table4.** Theta frequency analysis across all task epochs : no significant differences between task phases

| frequency (Hz) | #LE46 | #LE82 | #LE83 | #LE84 | #LE87 |
| --- | --- | --- | --- | --- | --- |
| retrieval | 7,81 | 8,06 | 8,02 | 8,81 | 8,15 |
| study | 8,29 | 8,04 | 8,1 | 8,01 | 8,37 |
| baseline | 8,33 | 8,17 | 8,21 | 8,12 | 8,25 |

**Supplementary table5.** Theta amplitudes are higher during retrieval but no difference between different trial types.

Mann-Witney U-test
| p-values | #46 | #82 | #83 | #84 | #87 |
| --- | --- | --- | --- | --- | --- |
| **amp retrieval > study** | 0,0026* | 0,0043* | 1,86E-07* | 5,6E-04* | 0,0002* |
| **amp retrieval>baseline** | 0,0001* | 0,0031* | 0,0002* | 0,0001* | 0,0107* |
| **amp study > baseline** | 0,2622 | 0,2141 | 0,02 | 0,089 | 0,1993 |
| **repeated vs. Non-repeated** | 0,48 | 6,73E-04* | 0,23 | 0,23 | 0,068 |
| **pre vs. In-trial** | 0,465 | 0,028* | 0,012* | 0,174 | 0,108 |

Abbreviations: amp: amplitudes; pre: pre-trial.
All theta measured at most distCA1
\*significant, Bonferroni corrected, $p < 0,05/3 = 0,0167$

**Supplementary Table6.** Speed of naive and trained animals and comparisons

| **speed(cm/sec)** |  |  |  |  |  |  |
| --- | --- | --- | --- | --- | --- | --- |
| **trained rats** | **baseline** | **study** | **test** | **freerun** | **naive rats** | **baseline** |
| LE46 | 2,8978 | 4,36 | 4,18 | - | LE1063 | 0,2114 |
| LE82 | 0,1455 | 4,34 | 5,05 | - | LE1080 | 1,9265 |
| LE83 | 0,5315 | 4,04 | 5,16 | 13,92 | LE1083 | 2,3689 |
| LE84 | 0,9305 | 4,5 | 4,09 | 22,5 | LE1085 | 0,0195 |
| LE87 | 2,2611 | 4,48 | 4,95 | - | LE1086 | 0,4334 |

**2-sample t-test**
| **group stats** | **t** | **p** |
| --- | --- | --- |
| freerun vs. Test | 5,8353 | 0,0021 |
| freerun vs study | 6,0535 | 0,0018 |
| freerun vs baseline | 6,88 | 9,93E-04 |

**Supplementary table7.** Comparing amplitudes of theta cycles between distCA1 and proxCA1, Mann-Witney U-test

| **trained** |  | **#46** | **#82** | **#83** | **#84** | **#87** | **#86** |
| --- | --- | --- | --- | --- | --- | --- | --- |
| p value | retrieval | 1,32E-20 | 6,95E-11 | 2,77E-20 | 1,80E-21 | 7,90E-16 | - |
|  | study | 2,97E-18 | 1,56E-33 | 6,62E-19 | 4,51E-09 | 2,70E-03 | - |
|  | baseline | 3,59E-34 | 9,42E-15 | 2,47E-46 | 5,05E-19 | 1,75E-07 | - |
|  | freerun |  |  | 1,63E-37 | 9,56E-31 |  | 2,43E-23 |
| **naïve** |  | **#1063** | **#1080** | **#1083** | **#1085** | **#1086** |  |
| p value | baseline | 7,7E-09(p>d) | 1,32E-32 (p>d) | 0,43 | 0,3 | 1,46E-4(d>p) |  |
|  | freerun | 8,64E-68(p>d) | 5,58E-74(p>d) | 3,50E-01 | 4,22E-44(d>p) | 5,49E-4(d>p) |  |

Abbreviations: p>d or d>p: proximal CA1>distal CA1 or vice versa
In trained rats all d>p.

**Supplementary Table8.** Comparing the theta amplitudes between in distal or proximal CA1

| **KS test** | **baseline** |  | **study** |  | **test** |  |
| --- | --- | --- | --- | --- | --- | --- |
| **trained rats** | D-value | stats(p) | D-value | stats(p) | D-value | stats(p) |
| **LE46** | 0,59 | 7,01E-37 | 0,64 | 5,10E-22 | 0,72 | 2,68E-25 |
| **LE82** | 0,27 | 1,19E-12 | 0,63 | 1,13E-36 | 0,35 | 1,12E-10 |
| **LE83** | 0,33 | 2,15E-42 | 0,28 | 2,89E-15 | 0,29 | 2,78E-15 |
| **LE84** | 0,45 | 6,55E-21 | 0,42 | 6,17E-09 | 0,63 | 1,11E-16 |
| **LE87** | 0,21 | 7,05E-05 | 0,23 | 5,80E-03 | 0,52 | 2,23E-13 |

| **naive rats** | D-value | stats(p) |
| --- | --- | --- |
| **LE1063** | 0,17 | 1,16E-05 (p>d) |
| **LE1080** | 0,23 | 1,26E-09 (p>d) |
| **LE1083** | 0,39 | 4,36E-27 (p>d) |
| **LE1085** | 0,07 | 2,20E-01 (d>p) |
| **LE1086** | 0,1 | 2,00E-03 (d>p) |

Kolmogorov-Smirnov (KS) test for baseline: trained vs naive rats: p=0,0079
p>d: theta amplitudes in proxCA1 are larger than in distCA1
d>p: theta amplitudes in distCA1 are larger than in proxCA1
F(X)=observed cumulative frequency distribution of a random sample of n observations
Friedman's test for trained animals, baseline vs. Study vs. Test p=0,2466
Wilcoxon ranksum test for comparing trained vs. Naive animals during baseline: p=0,0079

**Supplementary Table9.** Comparing theta amplitudes and speed between freerun and task phases during the delayed non-matching to odor task 2 tailed Mann-Whitney U-test

|  | #LE83 |  | #LE84 |  |
| --- | --- | --- | --- | --- |
| p value | theta amp | speed | theta amp | speed |
| **freerun vs.** |  |  |  |  |
| **Retrieval** | 8.5E-05 | 1,37E-10 | 0,3333 | <0.001 |
| **freerun vs.** |  |  |  |  |
| **study** | 3,07E-26 | 7,33E-12 | 9,10E-15 | <0.001 |
| **freerun vs.** |  |  |  |  |
| **baseline** | 4,19E-16 | <0.001 | 3,57E-14 | <0.001 |

Abbreviation amp: amplitudes.

