## Supplementary material for "Phase-locking of hippocampal CA3 neurons to distal CA1 theta oscillations selectively predicts memory performance"

### Supplementary results

#### SVM analysis using firing patterns of individual hippocampal subregions and 'firing rate different' neurons

To examine whether neuronal firing patterns in individual hippocampal subregions may discriminate between 'repeated' and 'non-repeated' odor trials we performed SVM analyses using the population firing activities in each hippocampal subregions (distCA1, proxCA1, distCA1 and proxCA3 separately). The SVM performances of neurons in the ltater brain areas did not differ from chance level (all p>0.06, Supp. Fig. 2A, Supp. Table 1), suggesting that neuronal firing patterns in these regions cannot predict memory performance.

To examine whether combining the firing activities of the neurons that fires differentially for repeated vs. non-repeated odor trials (i.e. the 'firing rate different' neurons) can predict memory performance, we also performed SVM analysis with this population of neurons. Combining the firing rates of these neurons could predict memory performance (SVM performance: 61%, t(4)=11, p=1.94e-04, one-tailed t-test to chance level; Supp. Table1, Supp. Fig. 2B) but to a lower extent than combining the firing rates of 'effective' neurons that were comparable in number (9 vs 8; comparison to SVM performance of effective neurons (85% correct); t(8)=5.5799, $p=5.22x10^{-4}$, 2-sample t-test).

#### No general linear dependency between theta amplitudes and speed during the delayed non-matching to odor task

To test whether there was a dependency between theta amplitudes and the speed of animals during the study or the test phase of the delayed non-matching to odor task, we extracted the local field potential, applied cycle-by-cycle analysis at theta frequency (6-12Hz) and aligned the output of this analysis and the instantaneous speed of the animals over the 2 sec of each trial (10 study trials and 20 test trials, one example test trial is displayed in Supp. Fig8A). For a direct comparison, the theta amplitude of each cycle was plotted as a function of speed estimated by averaging the speed during a 80ms time-window, centering at the middle of the peak and the trough of the theta cycle. A linear regression and a least square approach were applied to test for the dependency between amplitudes and speed. For each animal, the study- and test-phase are fit with an independent model. As shown in Supp. Fig. 8B-F, in the majority of the cases (8/10), the slopes of the linear fit were flat (but for the study phase of LE46 and LE84, p<0.05) showing, overall, no rigorous dependency between theta amplitudes and speed during the present non-spatial task. Note that the average speed of animals in this task is very low (<5cm/sec) and comparable to the range of speeds excluded from studies reporting a strong dependency between theta power and animals' speed (Kaefer et al., 2020; Stella et al., 2019)).

#### Freerun: speed and theta amplitudes

The average speed during freerun (i.e. when rats were given the opportunity to collect food pellets scattered in an open field; 18.26+-6.12cm/sec) of trained animals is higher than that in all other phases of the task (all Ps< 0.0021, Bonferroni corrected, Supp. Table 9).

In terms of theta amplitudes, in one rat (LE83), theta amplitudes were higher during freerun than all phases of the task (all ps $< 0.001$) while they were comparable between retrieval and freerun in the second rat (LE84, and significantly higher than study and baseline: freerun: vs retrieval: p=0.3; vs study or baseline: all ps<0.001, Supp. Table 9). Hence, no specific patterns could be found when comparing theta oscillation amplitudes at retrieval and freerun (Mann Whitney U-tests, Bonferroni corrected; see Supp. Table 9).

The proximodistal difference in theta amplitudes observed at test, study and baseline in trained animals was also observable in all trained rats which performed the freerun test (3/3; all $p<2.43\times10^{-23}$, Kolmogorov–Smirnov test, Bonferroni corrected, Supp. Table 7; of note, no study/test data for 1 rat). A distribution of one representative animal is shown in Fig. 4D. In contrast, no clear patterns could be identified in naïve animals (2 rats with amplitudes higher at proximal than distal CA1, 2 rats higher in distal than proximal CA1 and no difference for the last rat, Supp. Table 7, Supp. Fig. 9).

**Effect of CA1 theta power on CA3 spike-CA1 theta phase locking information**

To assess whether differences in phase-locking to distal versus proximal CA1 could be explained by a higher theta power in distal CA1, we added the normalized theta power (z-score transformed) as one additional feature to the theta phase matrix and report that adding theta amplitudes does not significantly alter SVM performance at the group level (2-sample t-test, t(8)=0,4851,p=0,6406; see also the table below forh individual SVM performances).

| SVM perf (%) | Animal #46 | #82 | #83 | #84 | #87 | average |
|---|---|---|---|---|---|---|
| Use theta phase only | 80 | 75 | 90 | 85 | 80 | 82 |
| Adding theta power | 50 | 75 | 95 | 90 | 80 | 78 |

**Comparing SVM performances of CA3 phase-discriminating neurons phase-locked to repeated and non-repeated stimuli to those of CA3 phase-discriminating neurons phase-locked to both types of stimulus AND to only one type of stimulus**

Among all the CA3 phase-discriminating neurons (69 neurons in total), 20 are significantly phase-locked to one odor type while 49 were phase-locked to different angles of distCA1 theta for the two stimulus types. To test whether the neurons which are phase-locked to only one stimulus type also contribute significantly to the SVM performance, we performed a separate SVM analysis using only the 49 neurons phase-locked to different angles of distCA1 for both stimulus conditions. Results show that SVM performance in this case is lower than that using all 69 phase-discriminating neurons (t(8)=3.9, p=0.0045, 2-sample t-test) and is reduced by about 15% for all animals (see the table below), suggesting that neurons phase-locked to only one condition also partially contribute to SVM performance.

| SVM perf (%) | Animal #46 | #82 | #83 | #84 | #87 | average |
|---|---|---|---|---|---|---|
| All phase discriminative neurons | 80 | 75 | 90 | 85 | 80 | 82 |
| Only neurons phase-locked during both conditions | 70 | 65 | 75 | 60 | 55 | 65 |

Kaefer, K., Nardin, M., Blahna, K., and Csicsvari, J. (2020). Replay of Behavioral Sequences in the Medial Prefrontal Cortex during Rule Switching. Neuron *106*, 154-165 e156. 10.1016/j.neuron.2020.01.015.

Stella, F., Baracskay, P., O'Neill, J., and Csicsvari, J. (2019). Hippocampal Reactivation of Random Trajectories Resembling Brownian Diffusion. Neuron *102*, 450-461 e457. 10.1016/j.neuron.2019.01.052.
